# Structure of PCPE-2 in complex with the BMP-1 metalloprotease reveals the molecular basis of its mechanism of inhibition

**DOI:** 10.64898/2026.09.22.752281

**Authors:** Julien Bauer, Priscillia Lagoutte, Jonathan Vaneyck, Valeria Calvaresi, Emmanuel Bettler, Mireille Dumoulin, Catherine Moali, Loïc Carrique, Sandrine Vadon-Le Goff

**Author notes:** Corresponding authors: Catherine Moali, Loïc Carrique, Sandrine Vadon-Le Goff. These authors contributed equally to the work (Julien Bauer, Priscillia Lagoutte).

## Abstract

PCPE-2 (procollagen C-proteinase enhancer-2) is an extracellular glycoprotein playing dual functions in the regulation of BMP-1/tolloid-like proteinases. Like its homologue PCPE-1, PCPE-2 can enhance the proteolytic maturation of fibrillar procollagens, but it also acts as a potent and specific inhibitor of BMP-1 through formation of a high-affinity complex with the protease. Here, we investigated the molecular basis of this inhibitory interaction using complementary biochemical, biophysical and structural approaches. We show that the two CUB domains of PCPE-2 must be covalently linked for efficient BMP-1 binding and inhibition, consistent with cooperative engagement of the protease. Cryo-electron microscopy combined with density-guided structural modelling and hydrogen/deuterium-exchange mass spectrometry supports a bipartite interaction complex in which the CUB1 domain of PCPE-2 engages the catalytic domain of BMP-1, while the CUB2 domains of both proteins interact together. The interaction involves the Ca²⁺-binding surfaces of the PCPE-2 CUB domains that also participate in procollagen recognition, providing a structural framework for understanding the mutually exclusive interactions of PCPE-2 with BMP-1 and procollagen. Allosteric inhibition is supported by the binding of PCPE-2 CUB1 opposite to the active site cleft, leading to the shielding of the residues surrounding the S1’ pocket, as observed by HDX-MS. Together, these findings define the architecture of the inhibitory PCPE-2/BMP-1 complex and provide the molecular basis for understanding the distinct regulatory activities of the two procollagen C-proteinase enhancers.

## Introduction

The BMP-1/tolloid-like proteinases (BTPs) form a small family of four secreted metalloproteases that play essential roles in development, tissue homeostasis, and extracellular matrix (ECM) organization^1,2^. These evolutionarily conserved enzymes, which include BMP-1 (bone morphogenetic protein-1), mammalian tolloid (mTLD; a splice variant of BMP-1), and mammalian tolloid-like 1 and 2 (mTLL-1, mTLL-2), share a multi-domain architecture comprising a catalytic (cat) astacin-like metalloprotease domain followed by CUB and EGF-like domains. Beyond their well-established functions in ECM assembly through the proteolytic maturation of several fibrillar and non-fibrillar collagens, proteoglycans, laminins, mineralization factors and lysyl oxidases, BTPs also modulate signalling pathways by processing growth factor precursors or regulators, angiogenic factors, adipokines and hormones^3,4^. Thus, they are key players in morphogenesis^5^, bone homeostasis^6,7^, muscle growth^8^, lipid metabolism^9^ or neurogenesis^10^. Altogether, this pleiotropy underscores their central position at the interface between ECM assembly and cell signalling, with important contributions to tissue homeostasis and tissue repair^11^, and implications in pathological processes such as fibrosis^12,13^ or tumour progression^14,15^.

The broad biological impact of BTPs is expected to require a tight regulation of their proteolytic activity. Several inhibitors^1,16^ have been reported but evidence for a tight-binding and specific endogenous inhibitor of human BTPs was lacking until recently. For example, BTPs are inhibited by α2-macroglobulin through the well-described protease-trap mechanism induced upon cleavage of a bait region^17^. However, this mechanism is common to most proteases and not specific to BTPs. Also, the extracellular protein called Sizzled found in *Xenopus*, chicken or zebrafish, inhibits BTPs from these organisms but is absent in mammals, and reports describing human sFRP-2 (secreted Frizzled Related Protein-2) as an homologous inhibitor have not been consistently reproduced across laboratories^1,2^. Another such controversial inhibitor is BMP-4. Initial studies reported that BMP-4 interacts with the prodomain of BMP-1 but not with its mature catalytic form, suggesting that the prodomain acts as a BMP-4 antagonist^18^. However, subsequent studies demonstrated that BMP-4 is able to bind the mature form of *Xenopus* BMP-1 through its CUB1 and CUB2 domains^19^, but this has never been confirmed in mammals.

In contrast, BTP activity is known to be efficiently upregulated by extracellular modulators that enhance the cleavage of specific substrates^1,2^, such as TSG (twisted gastrulation) for chordin, periostin for lysyl oxidase, WFIKKN for myostatin or PCPE-1 (procollagen C-proteinase enhancer-1) for fibrillar procollagens. PCPE-1 is the most well-studied of these substrate-specific regulators and was shown to act by binding the procollagen substrate to induce a conformational change and facilitate its proteolytic processing by BTPs^20^.

PCPE-2, a homologue of PCPE-1, was also reported to enhance procollagen cleavage by BMP-1, supporting the view that the two PCPEs were functionally interchangeable^21^. However, a recent study^22^ demonstrated that PCPE-2 also acts as a direct and potent inhibitor of BMP-1, affecting all substrates, through the formation of a tight-binding complex. This unexpected activity is mediated by the astacin-CUB1-CUB2 region of BMP-1 (Fig. 1A; described as [catCUB1CUB2] hereafter) and the CUB1-CUB2 domains of PCPE-2 (Fig. 1B; described as CUB1CUB2 hereafter) and efficiently prevents BMP-1 proteolytic activity with a K_i_ in the low nanomolar range^22^. Despite its high homology with PCPE-2 (43% sequence identity and high structural similarity^23^), PCPE-1 neither binds nor inhibits BMP-1 in the same concentration range^22^. Thus, the inhibitory capacity is unique to PCPE-2 and this protein appears to be the missing specific endogenous inhibitor of mammalian BTPs.

**Figure 1:**
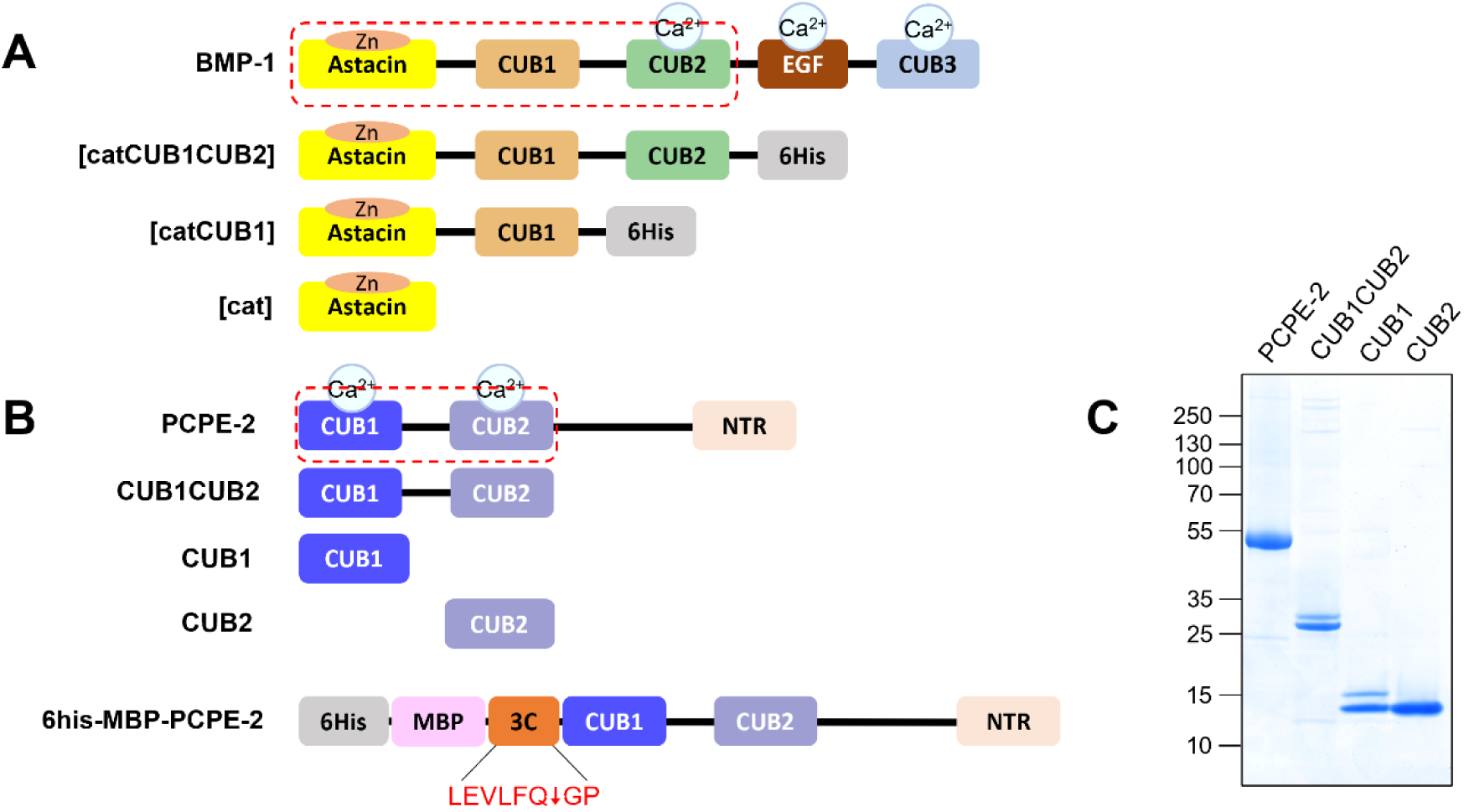
Constructs used in this study. **A** Domain structure of BMP-1 and its deletion mutants [catCUB1CUB2], [catCUB1], [cat]; minimal domains required for inhibition by PCPE-2 outlined with a red dotted box. **B.** Domain structure of PCPE-2 and of the different constructs used in the study; minimal domains of PCPE-2 necessary to inhibit BMP-1 also outlined in a red dotted box. The construct 6His-MBP-PCPE-2 was used to produce PCPE-2 and similar constructs were designed to obtain the CUB domains. **C.** SDS-PAGE analysis of PCPE-2 and its domains after purification (non-reducing conditions, 4-12% gel). The double bands visible in CUB1 and CUB1CUB2 originate from the heterogeneous O-glycosylation pattern of CUB1^22^.

The molecular basis for this striking functional divergence between PCPE-1 and PCPE-2 remains unclear. Understanding the structural and biochemical principles that govern the selective recognition of BMP-1 by PCPE-2 is critical, as it may reveal a new mechanism of BMP-1 regulation and allow the design of therapeutic modulators targeting ECM aberrant organization. To address these questions, we used an integrative approach combining biochemistry, molecular modelling, cryo-electron microscopy (cryo-EM), and hydrogen/deuterium-exchange mass spectrometry (HDX-MS). Our findings uncover the structural determinants of PCPE-2 specificity and provide broader insights into the selective regulation of BTP proteinases.

## Results

### The CUB domains of PCPE-2 need to be linked to interact with BMP-1 or to inhibit its activity

To define the respective contributions of the PCPE-2 CUB1 and CUB2 domains to BMP-1 binding and inhibition, we produced full-length PCPE-2, CUB1CUB2, and the isolated CUB1 and CUB2 domains (Fig. 1B). Because previous expression strategies for the PCPE-2 constructs yielded unstable preparations and did not allow the production of isolated CUB2, we used an MBP-fusion strategy incorporating an HRV 3C protease cleavage site. Following proteolytic removal of the N-terminal 6His-MBP tag, all four PCPE-2 constructs could be purified to homogeneity, with yields ranging from 1.2 to 4.8 mg/L (Fig. 1C). NanoDSF analysis confirmed that the isolated CUB domains were properly folded and displayed similar thermal stability (Fig. S1), indicating that differences in their interaction with BMP-1 would unlikely result from differences in protein stability. In parallel, we used a series of BMP-1 constructs^22^ comprising its catalytic domain and increasing numbers of CUB domains (BMP-1 [cat], BMP-1 [catCUB1] and BMP-1 [catCUB1CUB2]; Fig. 1A), with BMP-1 [catCUB1CUB2] corresponding to the minimal BMP-1 region previously shown to support PCPE-2 binding and inhibition.

We compared the interaction of PCPE-2 constructs with full-length BMP-1 using surface plasmon resonance ((SPR), Fig. 2A). CUB1 showed only very weak binding to BMP-1, whereas CUB2 exhibited a stronger binding, with an equilibrium dissociation constant around 400 nM (Fig. 2B), markedly higher than that reported for CUB1CUB2 (28 nM)^22^. CUB2 also displayed faster association and dissociation rates than CUB1CUB2, resulting in the lower affinity for BMP-1 (Fig. S2). In addition, co-injection of the isolated CUB1 and CUB2 domains over BMP-1 produced a response that was similar to that of CUB2 alone and substantially lower than that obtained with the covalently linked CUB1CUB2 (Fig. 2A), indicating that the enhanced binding of CUB1CUB2 requires their physical linkage and cannot be reproduced simply by providing the two domains simultaneously.

**Figure 2:**
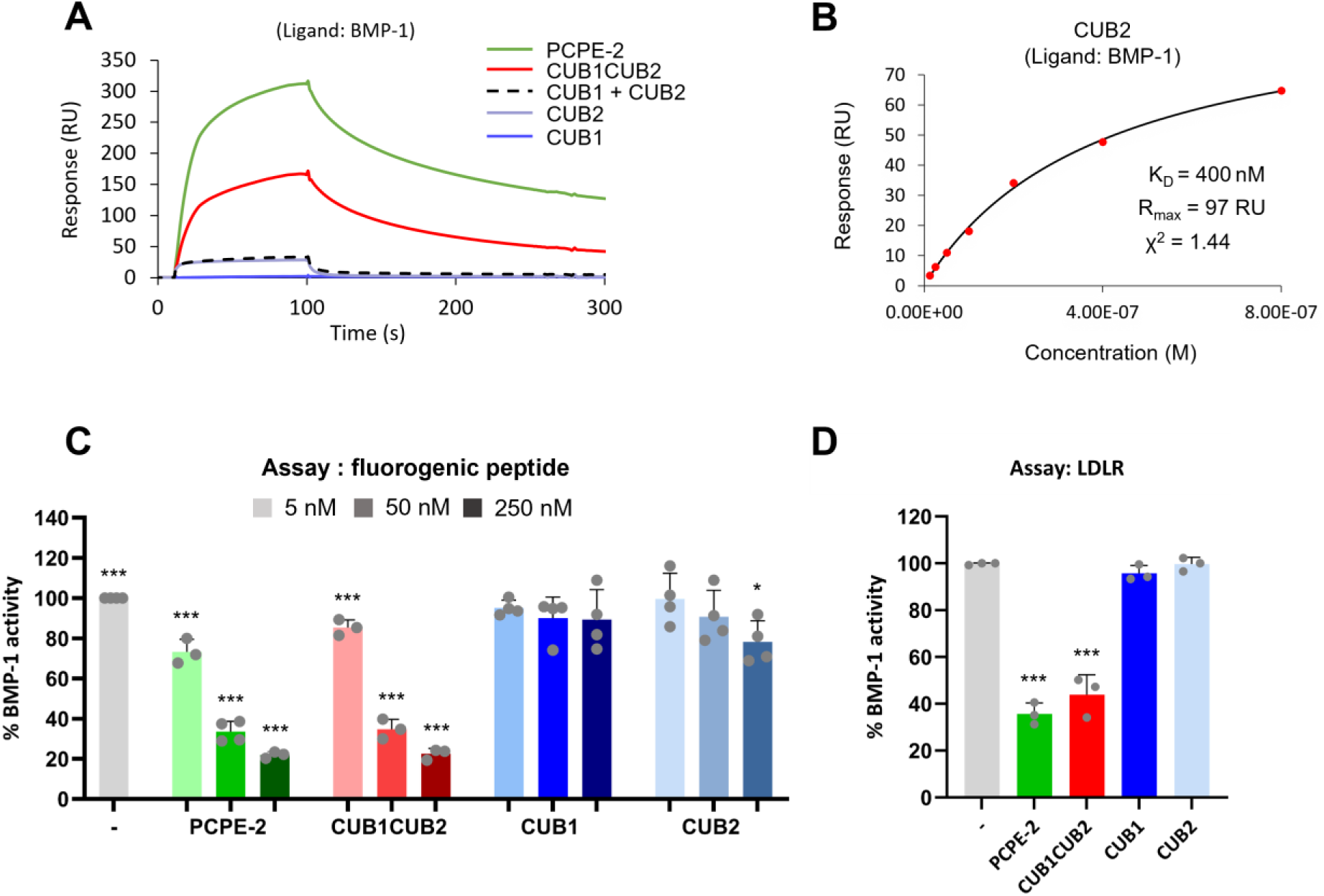
The CUB domains of PCPE-2 need to be linked to interact and inhibit BMP-1. **A.** Comparison of the binding of 100 nM PCPE-2 and its domains (CUB1CUB2, CUB1, CUB2, equimolar mixture of CUB1 and CUB2) over immobilized BMP-1 (1640 RU). **B.** Fit (steady state model) of the SPR response at the end of injection, obtained when increasing concentrations of CUB2 (12.5 nM – 800 nM) were injected over immobilized BMP-1 (1640 RU). **C.** Quantification of BMP-1 activity with the fluorogenic peptide Mca-YVADAPK(Dnp)OH in the presence of increasing concentrations of PCPE-2 and its deletion mutants. BMP-1 concentration in the experiment was 7 nM. Means ± SD of n=3 to 4 independent experiments run in duplicate. **D.** Bar charts showing the effect of PCPE-2 and its deletion mutants (388 nM) on the cleavage of LDLR ectodomain (388 nM) by BMP-1 (13 nM). Means ± SD of n = 3 independent experiments performed in duplicate. Uncropped gels available in Supplementary Fig. S3. Statistical significance (comparison with the BMP-1 alone condition): p-value ≤ 0.05 (*), p-value ≤ 0.001 (***).

A similar trend was observed for the inhibition of BMP-1 proteolytic activity. While PCPE-2 and CUB1CUB2 inhibited the cleavage of a fluorogenic peptide by BMP-1 in a concentration-dependent manner, consistent with our previous study^22^, CUB1 had no detectable inhibitory effect and CUB2 showed only a slight but significant inhibition at the highest concentration tested (Fig. 2C). The cleavage of the LDLR ectodomain, a physiological substrate of BMP-1^24^ was then investigated. Under the condition of the experiment, full-length PCPE-2 and CUB1CUB2 strongly inhibited LDLR cleavage, whereas neither isolated CUB domain produced significant inhibition (Fig. 2D). Altogether, these results demonstrate that physical linkage of the PCPE-2 CUB1 and CUB2 domains is required for efficient BMP-1 binding and inhibition.

### Development of BMP-1-targeting VHHs to better localize the PCPE-2 interaction site

We next sought to better define the regions of BMP-1 involved in the interaction. Previous studies have established that the BMP-1 [catCUB1CUB2] domains were necessary for the formation of the PCPE-2/BMP-1 complex^22^. Attempts to produce the isolated BMP-1 CUB domains, including as MBP fusion proteins, were unsuccessful. Therefore, we adopted an alternative strategy and generated VHHs (variable domains of heavy-chain only antibodies; also known as nano-antibodies or nanobodies®^25^) targeting BMP-1.

An immune library was created from the blood of an alpaca immunized with BMP-1 [catCUB1CUB2] following established protocols^26^ (Fig. 3A). VHHs were selected through three rounds of phage display, using BMP1 as a bait to enrich the library in phage particles displaying VHHs specific of BMP1. A small-molecule inhibitor (UK383,367), which binds into the BMP-1 active site, was added to prevent potential BMP-1-mediated cleavage of the VHHs. Ninety of the clones obtained after rounds 2 and 3 of the panning were randomly selected and screened by indirect ELISA to detect the presence of anti-BMP-1 VHHs. Fifteen clones gave a positive signal and their VHH gene was sequenced. Three VHHs belonging to three different families according to the sequence of their CDR3 were identified: NbB198, NbB207 and NbB220. The length of their CDR3 is comprised between with 16, 13 and 18 residues respectively. Interestingly, NbB207 and NbB220 have the same CDR1 and CDR2 while their CDR3 greatly differ (Fig. S4).

**Figure 3:**
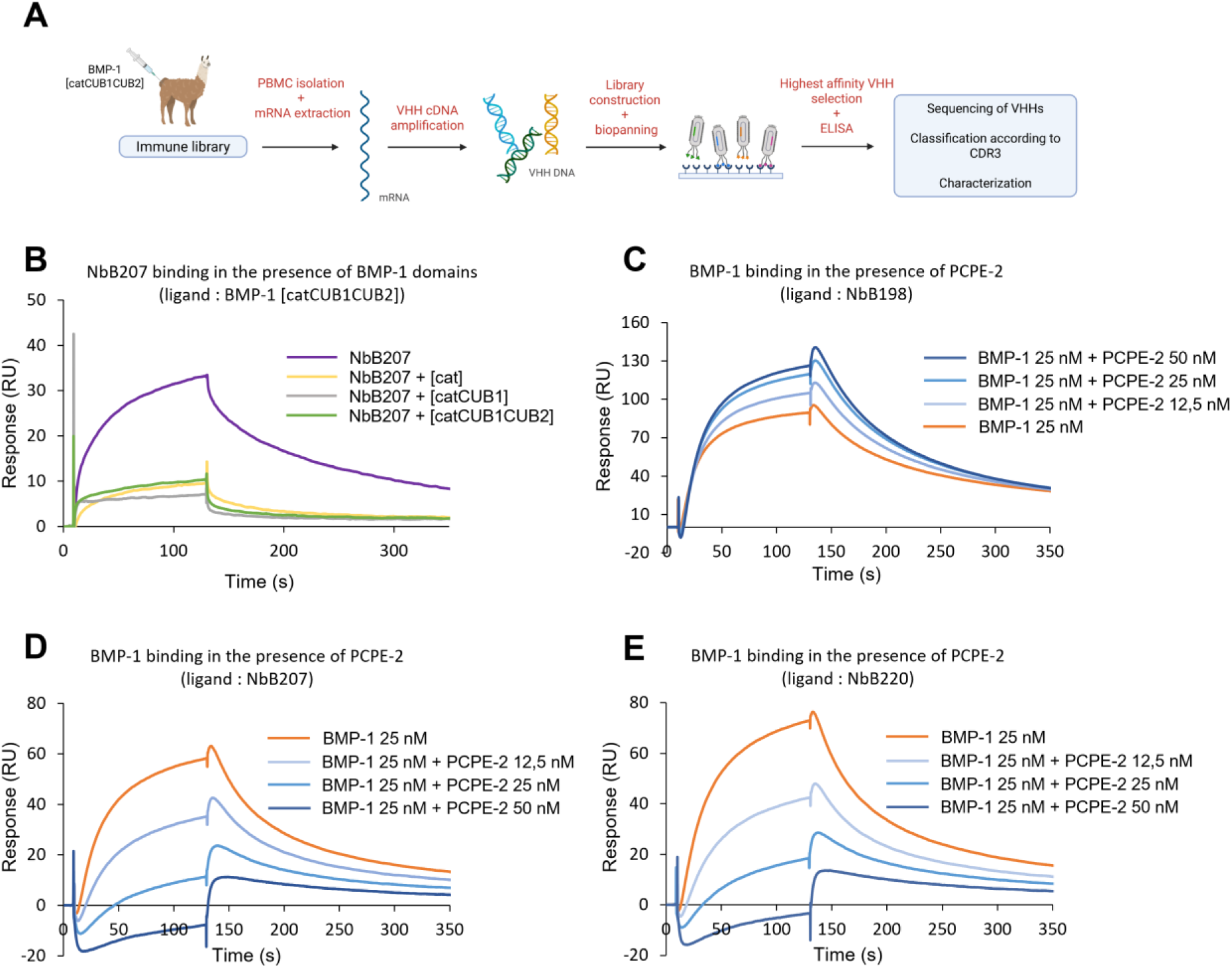
Selection of VHHs against BMP-1 and characterization of their interaction profiles. **A.** Schematic workflow of the generation and selection of the VHHs targeting BMP-1 [catCUB1CUB2]. **B.** Competition experiment in which 100 nM of NbB207 were co-injected with 100 nM of BMP-1 truncated variants ([cat], [catCUB1], [catCUB1CUB2]). The immobilized protein was BMP-1 [catCUB1CUB2] (397 RU). **C.** Competition experiment consisting of an injection of a mixture of 25 nM of BMP-1 with increasing concentrations of PCPE-2 (12.5 nM, 25 nM, 50 nM). The immobilized protein was an anti-His antibody (13605 RU) for the capture of NbB198 (48 RU). **D.** Same as in (C) but with NbB207 (53 RU). **D.** Same as in (C) but with NbB220 (53 RU).

The three VHHs were produced in the periplasm of *E. coli* and purified as described^27^. All appeared as monomers in following SEC analysis. Their affinity for BMP-1 was evaluated using SPR, showing that they all have high affinities for the target, with K_D_s ranging from 9 nM (NbB198) to 20 nM (NbB207) and 24 nM (NbB220) (Fig. S5).

We then mapped the BMP-1 regions recognized by the VHHs using competition experiments with the truncated versions of BMP-1 (Fig. 1A, 3B and S6). Co-injection of each VHH with any of the three BMP-1 constructs strongly reduced VHH binding to immobilized BMP-1 [catCUB1CUB2], with comparable competition observed for all three constructs (Fig. 3B and Fig. S6). Since the catalytic domain is the only region shared by these constructs, these results indicate that all three VHHs recognize epitopes located within, or primarily determined by, the catalytic domain of BMP-1. Consistent with binding outside the active-site cleft, NbB198 and NbB220 had no detectable effect on BMP-1 activity, whereas NbB207 produced only modest inhibition (<20%) even when present at a 143-fold molar excess over the protease (Fig. S7). Pairwise co-injection experiments further showed that the three VHHs can bind BMP-1 simultaneously, indicating that they recognize non-overlapping epitopes within the catalytic domain (Fig. S8).

We next investigated whether there was a competition between PCPE-2 and the VHHs for BMP-1 binding. Co-injection of BMP-1 and increasing concentrations of PCPE-2 over the VHHs actually gave different results depending on the VHH (Fig. 3C-E). While BMP-1 binding to NbB207 or NbB220 was strongly reduced in the presence of PCPE-2, its interaction with NbB198 was much less affected. These results suggest that PCPE-2 shares overlapping epitopes with NbB207 or NbB220 but not with NbB198, and further support the fact that the catalytic domain of BMP-1 is required for PCPE-2 binding, as previously proposed^22^. Unfortunately, the requirement for additional BMP-1 domains could not be probed as no VHH against BMP-1 CUB domains could be obtained.

### Cryo-EM analysis of the PCPE-2/BMP-1 complex

We next sought to determine the structure of the PCPE-2/BMP-1 complex using single-particle cryo-electron microscopy (cryo-EM). To limit protein denaturation and aggregation at the air–water interface during vitrification, the complex was PEGylated before grid preparation^28^. The complex formed by the two full-length proteins was successfully isolated by size exclusion chromatography (SEC) (Fig. S9) and cryo-EM data were collected. The structure was determined to a nominal resolution of 3.64 Å (Fig.S10). Although the intrinsic flexibility and partial preferential orientation of the complex were limiting the interpretability of the map, we could unambiguously identify that the density was corresponding to the “minimal complex” composed of BMP-1 [catCUB1CUB2] and PCPE-2 CUB1CUB2 and that the other domains (NTR from PCPE-2 and EGF-CUB3 from BMP-1) were not visible. Further attempts to reduce preferred particle orientation on the grids or control the high flexibility of the proteins using the shorter versions of the partners were unsuccessful, as was the use of a megabody^29^ derived from B198 to form the PCPE-2/BMP-1/Megabody B198 complex (data not shown).

Due to the high structural similarity of the four CUB domains present in the complex, assignment of the partners in the density was very challenging. To go ahead, we generated an AlphaFold3^30,31^ model of the minimal BMP-1/PCPE-2 complex, including one Zn²⁺ and three Ca²⁺ ions (one for BMP-1 and two for PCPE-2) required for the structural integrity and activity of the proteins^32–36^. The individual domains were predicted with high confidence, as indicated by their pLDDT (predicted Local Distance Difference Test) scores (Fig. S11A), but their relative arrangement within the complex was poorly supported, with an interface predicted template modelling (ipTM) score of 0.15 only. Three independent predictions consistently produced similar arrangements with likewise low interface-confidence scores. In these models (Fig. 4A), the PCPE-2 CUB domains interacted predominantly with the catalytic domain of BMP-1, with no contribution from BMP-1 [CUB2], despite previous biochemical evidence showing that this domain is required for high-affinity complex formation^22^. In addition, fitting of the predicted complex into the cryo-EM density revealed than only the BMP-1 [catCUB1CUB2] could be accommodated within the map, whereas the predicted position of PCPE-2 was largely outside the reconstructed density (Fig. S11B).

**Figure 4:**
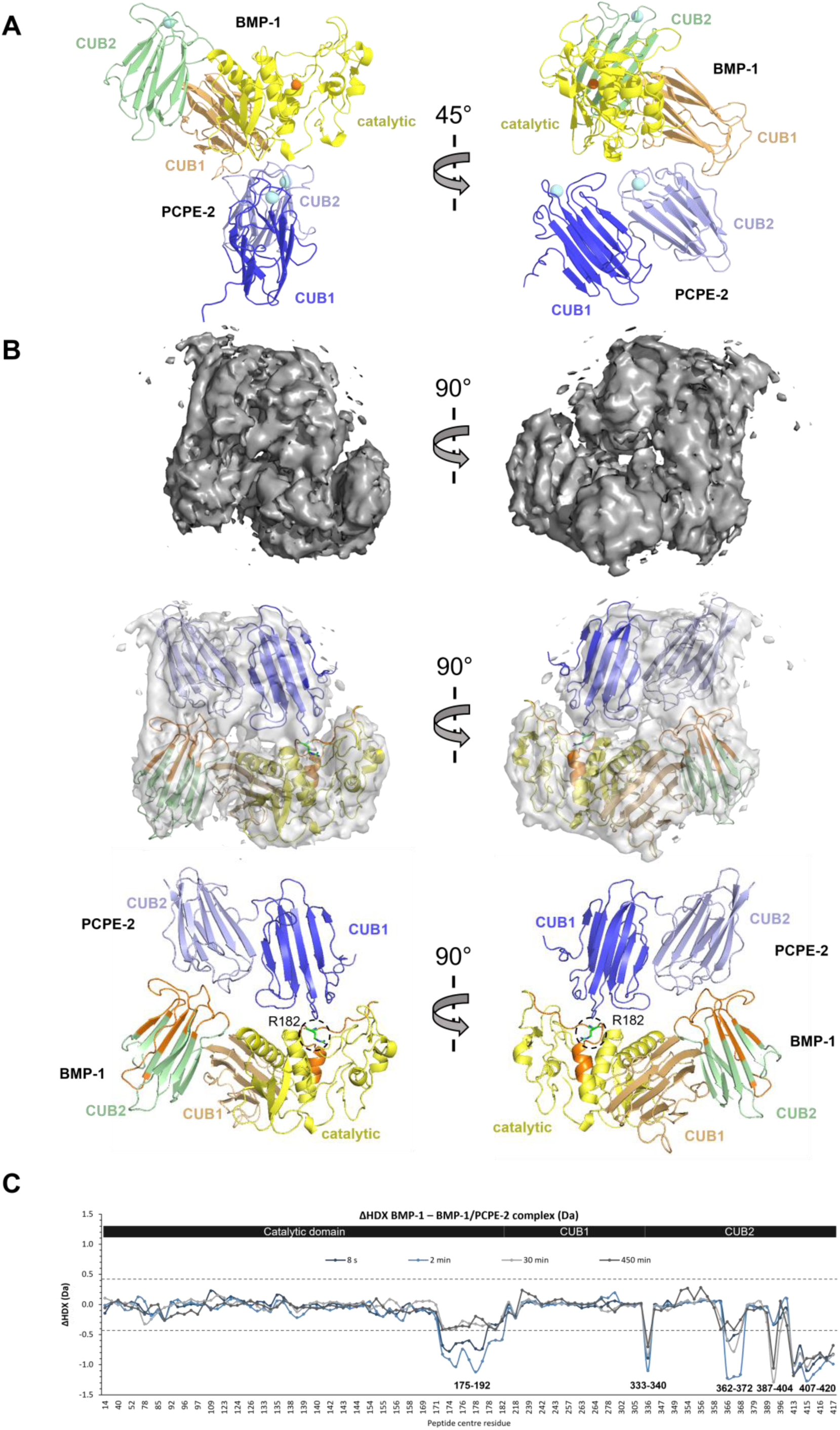
PCPE-2 interacts with the catalytic domain and the CUB2 domain of BMP-1. **A.** AlphaFold3 model of PCPE-2 CUB1CUB2 (blue) in complex with BMP-1 [catCUB1CUB2]. Ca^2+^ ions are represented in light blue and Zn^2+^ ion in orange. This model is representative of three independent predictions. **B.** Top, cryo-EM density map of the BMP-1 [catCUB1CUB2] – PCPE-2 CUB1CUB2 complex at 3.64 Å nominal resolution. Middle, CryoZeta structural model fitted into the corresponding cryo-EM density map. Bottom, CryoZeta structural model with BMP-1 [catCUB1CUB2] and PCPE-2 CUB1CUB2. Residues identified by HDX-MS as being involved in the interaction are highlighted in orange and the R182 that dictates protease substrate specificity is showed in green sticks and surrounded by dotted lines. BMP-1 is positioned in the same orientation as in the AlphaFold3 model. **C.** Plot illustrating the differences in HDX (ΔHDX) between BMP-1 alone and BMP-1 in complex with PCPE-2 across the time points studied. X-axis: peptides whose HDX was followed are ordered according to their center residue. Y-axis: difference in HDX (Da). The dashed gray lines represent the thresholds of significant differences in HDX.

In order to improve our modelling, we used CryoZeta^37^, a density-guided structure-prediction approach that integrates experimental cryo-EM density with diffusion-based structural modelling, generating models constrained by the experimental map. The best-scoring model showed good agreement with the cryo-EM density, with CCmask and CCbox values of 0.69 and 0.73, respectively, and a Model Reliability Score of 0.69. In contrast to the AlphaFold3 prediction, both BMP-1 and PCPE-2 were fully accommodated within the experimental density (Fig. 4B, same orientation as in Fig. 4A for BMP-1). Structural superposition of the crystallographic structure of the catalytic domain of BMP-1 (the only domain for which structural data is available, 6BSM^38^) with the corresponding domains from the AlphaFold3 and CryoZeta models yielded root mean square deviation (RMSD) values of 0.3 Å and 0.5 Å, respectively, indicating that both approaches accurately reproduced the experimentally determined structure of this domain.

The CryoZeta model suggested that the CUB1 domain of PCPE-2 interacts with the catalytic domain of BMP-1 while the CUB2 domains of PCPE-2 and BMP-1 interact with each other. PCPE-2 is arranged with its two CUB domains positioned side by side, with their calcium-binding regions facing towards BMP-1 (Fig. 4B). Consistent with the stronger contribution of PCPE-2 CUB2 observed in the binding experiments, the interface formed by the CUB2-[CUB2] interaction is larger than the CUB1–catalytic-domain interface (Fig. 4B), with buried interface areas of 611 Å^2^ and 410 Å^2^, respectively. This architecture therefore provides a structural rationale for the weak BMP-1 binding retained by isolated CUB2 and for the substantially higher affinity achieved when CUB1 and CUB2 are covalently linked (Fig. 2A).

### Validation of the complex interface by HDX-MS

To experimentally assess the interaction architecture suggested by the CryoZeta model, we performed differential hydrogen/deuterium-exchange mass spectrometry (HDX-MS) using the minimal interacting constructs BMP-1 [catCUB1CUB2] and PCPE-2 CUB1CUB2. HDX-MS measures the peptide-level deuterium uptake of proteins exposed to a deuterated buffer for different time periods, allowing assessment of their local structural dynamics under different states^39^. We followed the HDX of 111 peptides of BMP-1, spanning 84.1% of the sequence and including three N-glycosylation sites (N22, N212 and N243; Fig. S12–S16 and Table S1). Comparison of BMP-1 alone and in complex with PCPE-2 revealed significant decreases in deuterium uptake in both the catalytic and CUB2 domains of BMP-1 (Fig. 4B-C). Within the catalytic domain, marked protection was observed for peptides spanning residues 175–192. This region (highlighted in orange in Fig. 4B) faces PCPE-2 CUB1 in the CryoZeta model, consistent with the predicted interaction between CUB1 and the BMP-1 catalytic domain. PCPE-2 binding also induced extensive protection within BMP-1 CUB2, including regions encompassing the loops connecting beta sheets β2/β3 (residues 333–340), β3/β4 (362–372), β5/β6 (387–404) and β7/β8 (407–420) (also highlighted in orange in Fig. 4B). Most of these regions map the predicted CUB2–CUB2 interface, providing independent experimental support for the involvement of BMP-1 CUB2 in complex formation. Notably, the protected region spanning residues 175–192 contains Arg182, a key determinant of BMP-1 substrate specificity that contributes to recognition of aspartate residues at the P1′ position of cleavage sites. In the CryoZeta model, PCPE-2 is positioned on the face of the catalytic domain opposite to the active-site cleft, indicating that inhibition is unlikely to result from direct steric occlusion of the active site. Instead, the altered HDX of the region surrounding Arg182 suggests that PCPE-2 binding affects the local environment or dynamics of a region involved in substrate recognition, supporting that allostery is a major component of PCPE-2 mechanism of inhibition.

## Discussion

Recent studies have established that PCPE-2 is a potent broad-spectrum inhibitor of BMP-1 activity, challenging its previous identification as an enhancer of procollagen processing, and raising questions about the functional divergence with the highly homologous PCPE-1 protein^22,23,40^. The present work investigated the molecular basis of the BMP-1/PCPE-2 interaction to understand how PCPE-2 modulates enzyme function at the structural and mechanistic levels.

One specificity of the PCPE-2/BMP-1 complex is the presence of multiple CUB domains in the two partners. CUB domains are widely distributed protein–protein interaction modules that are involved in a wide range of biological functions, especially in the extracellular environment^33^. It has been shown that Ca^2+^-binding CUB domains often interact with their binding partners through interactions between a basic side chain, from an arginine or a lysine residue, and the acidic residues that coordinate the Ca^2+^ ion within the CUB domain^41^. Interestingly, we evidenced that the CUB1 domain of BMP-1, the only CUB domain in the protease with no calcium binding site^36^, is not involved in the interaction with PCPE-2. Our data also indicate that PCPE-2 directly engages BMP-1 through a mechanism that requires the coordinated binding of its two CUB domains to the enzyme, as the binding of CUB2 alone remains weak and as CUB1 does not bind in the tested conditions. This mode of interaction is common in CUB-mediated protein-ligand binding, for which attachment of several CUB domains usually confers high-affinity binding through avidity effects, in spite of the low affinity of individual interactions^33,35,41^.

The PCPE-2/BMP-1 interaction adds another layer of complexity to CUB domain-containing proteins. Not only do PCPE-2 CUB domains need to cooperate to bind to BMP-1 but also the formation of the complex depends on the interaction between the CUB2 domains of both proteins. The integrative structural biology approach employed in this study, combining HDX-MS, cryo-EM, competitions with VHHs and molecular modelling, enabled us to define the domain interfaces. BMP-1 CUB2 domain binds to PCPE-2 CUB2 through interactions involving their calcium binding sites, while the catalytic domain of BMP-1 interacts with the CUB1 domain of PCPE-2. This position PCPE-2 opposite to the substrate-binding pocket, suggesting an allosteric inhibition mechanism leading to altered substrate recognition and/or catalytic efficiency. Along this line, HDX-MS shows that Arg 182 of BMP-1 which defines the S1’ pocket and dictates protease specificity is protected from deuterium exchange in the presence of PCPE-2, suggesting that it becomes less accessible to solvent and substrate. Allosteric inhibition is also supported by the previous finding that inhibition is never complete (∼ 75 % at the highest concentrations) with a short peptide substrate^22^, ruling out mechanisms involving pure competitive inhibition. With longer physiological substrates however, complete inhibition is observed^22^, possibly because PCPE-2 hinders proper binding of those bigger substrates to the catalytic and non-catalytic domains. Indeed, the CUB2 domain of BMP-1 was previously suggested to be needed for optimal proteolytic activity with procollagens and chordin^42,43^. These results reveal a unique mechanism of BMP-1 inhibition by PCPE-2. In comparison, Sizzled has been shown to require only the catalytic domain of BMP-1 for inhibition, not its CUB domains, and to inhibit the protease through a general mechanism involving the binding of a flexible loop inside the active site cleft of BMP-1^44^.

Furthermore, PCPE-2 has been previously shown to interact with procollagens in a manner similar to PCPE-1, involving the Ca^2+^ binding surfaces of its two CUB domains^22^. Our results indicate that these same regions of PCPE-2 are also involved in its interaction with BMP-1, suggesting an overlap between the BMP-1 and procollagen binding interfaces. This provides a structural explanation for the inability of PCPE-2 to bind BMP-1 and procollagen simultaneously^22^. The structural basis for the lack of PCPE-1 binding to BMP-1 remains to be elucidated, and will be addressed by careful comparison of the residues differing between PCPE’s CUB domains. However, this mode of action already highlights how highly similar domain architectures can achieve fundamentally different regulatory outcomes.

Finally, our study provides methodological insights into the challenges and opportunities of modern structural biology. The inability of AlphaFold3 to accurately predict the PCPE-2/BMP-1 complex, despite its success with many other protein-protein interactions, underscores the limitations of current AI-based approaches for certain classes of multi-domain, flexible protein complexes. The opportunity offered by tools such as CryoZeta (based on OpenFold3) to combine AI-based modelling and experimental data is perfectly demonstrated here. This has broader implications for structural studies of extracellular matrix proteins, which, like our BMP-1/PCPE-2 complex, often exhibit limitations in cryo-EM studies: flexibility, repetitions of similar domains, and preferred orientations.

## Material and Methods

### Cloning, expression and purification of proteins

The constructs containing PCPE-2 CUB1CUB2, CUB1 and CUB2 domains were amplified by PCR from pHLm-MBP-3C-CUB1CUB2-6His^22^ and inserted into the pHLm-MBP2 plasmid (with a N-terminal 6his tag; gift from Luca Jovine^45^; Addgene plasmid # 72344) between the NotI or NheI and XhoI restriction sites. Full-length PCPE-2 was amplified from pHLm-MBP-3C-PCPE2-6his and inserted into the pHL-mMBP-2 plasmid between the NotI and SmaI restriction sites. All constructs were checked by Sanger sequencing (MicroSynth) using ApE for visualisation^46^. The primers used for cloning are listed in Table S2.

6his-MBP-PCPE-2 and its deletion mutants were produced in HEK293-F cells (ThermoFisher Scientific, #R79007) grown in suspension in FreeStyle 293 expression medium (Gibco) using sterile flasks (Corning) placed on an orbital shaker platform (Eppendorf) rotating at 125 rpm and 37 °C with 8% CO_2_. On the day of transfection, cells were resuspended at a cell density of 1.0·10^6^ cells/mL in FreeStyle 293 expression medium. For transfection, plasmid DNA and PEI 25K transfection agent (Polysciences, filtered on 0.2 μm filters) were first diluted separately in 1/20 of the total culture volume of Opti-MEM medium (Gibco) and kept at room temperature for 5 min before mixing. The mixture was further incubated for 15 min before addition to the cells. Conditioned media were collected 3 days after transfection, centrifuged at 1,000 g and mixed with protease inhibitors (0.25 mM Pefabloc (Roth) and 2 mM N-ethylmaleimide (Merck)) and centrifuged again at 10,000 g.

6His-MBP-PCPE-2 was purified through the His-tag on Ni-Sepharose Excel (Cytiva). The cell supernatant was loaded onto the resin, pre-equilibrated in Buffer A (20 mM HEPES pH 7.4, 0.2 M NaCl). Non-specifically bound proteins were washed out with Buffer A containing 20 mM and 50 mM imidazole, and the protein of interest was eluted with 250 mM imidazole. Calcium was added to a final concentration of 5 mM to the protein solution before cleavage by His-tagged HRV 3C-protease. The latter was added at a ratio of 1/30 (w/w) and the mixture was incubated for 1 h at 21°C, then at 4°C overnight. Purification of the cleavage mixture was achieved on a Heparin Sepharose-6 Fast Flow column (Cytiva), equilibrated with buffer A containing 5 mM CaCl_2_. After washing, the protein was eluted with increasing concentrations of NaCl (0.5 M, 1 M and 2 M). The purified protein was diluted to a final concentration of 0.5 M NaCl and 0.1% βOG (n-Octyl-β-D-glucopyranoside) was added. Proteins were stored in 20 mM HEPES pH 7.4, 0.5 M NaCl, 5 mM CaCl_2_ and 0.1% βOG at −80 °C (after flash-freezing) until use.

6his-MBP-CUB1CUB2, -CUB1 and -CUB2 were purified and cleaved by HRV 3C-protease using the same procedure. Purification of the cleavage products was achieved on Ni-NTA Agarose (Qiagen). The resin was equilibrated with 20 mM HEPES pH 7.4, 0.5 M NaCl, 5 mM CaCl_2_ before the cleavage mixture was loaded and the flow-through containing the protein of interest was collected. βOG was added at a final concentration of 0.1% and the purified proteins were stored as above.

Full-length BMP-1 and its truncation mutants were produced and purified as described previously^22^. Biotinylated BMP-1 [catCUB1CUB2] was obtained through enzymatic labelling. The sequence of the [catCUB1CUB2] domains was cloned into pHLAvitag3^47^ between the EcoRI and KpnI restriction sites and expressed by transient transfection in 293-F cells. After purification on Ni-Sepharose Excel (Cytiva), avitagged [catCUB1CUB2] was buffer-exchanged to 50 mM bicine, 100 mM potassium glutamate pH 8.3 with zebaspin columns (ThermoFisher). Ten µM avitagged [catCUB1CUB2] were biotinylated using 700 nM GST-BirA^48^ (Biotin protein ligase) in 50 mM bicine, 100 mM potassium glutamate pH 8.3 supplemented with 10 mM ATP (Merck), 50 µM d-biotine (Merck) and 10 mM MgCl_2_ during 4 h at 30 °C. GST-BirA was eliminated on Glutathion Sepharose (Cytiva), and excess biotin was removed by desalting 2 times on a zebaspin column previously equilibrated with 20 mM HEPES pH 7.4, 0.5 M NaCl, 2.5 mM CaCl_2_, 0.1% βOG. The biotinylation level was evaluated by loading on Streptavidin Sepharose and analysing the flow through by SDS-PAGE, and was estimated to be superior to 95%.

### NanoDSF

Protein thermal stability and aggregation were assessed by nano differential scanning fluorimetry (nanoDSF) using a Prometheus Panta instrument (NanoTemper Technologies). Protein samples were loaded into standard-grade capillaries and subjected to a temperature ramp from 20 to 90°C at a heating rate of 1°C/min. Measurements were performed at a protein concentration of 0.5 mg/mL in Hepes 20mM, NaCl 0.5M, CaCl_2_ 5mM, βOG 0.1% pH 7.4. Data were acquired with PR.ThermControl version 2.3.1 and analysed with PR.Stability Analysis version 1.1.

### Selection and characterization of VHHs

Anti-BMP-1 VHHs were isolated from the immune phage library JOE3, generated from the blood of an alpaca immunized with BMP-1 [catCUB1CUB2] as previously described^26^. VHHs were selected through three rounds of solution-phase phage display panning against biotinylated, Avi-tagged BMP-1 [catCUB1CUB2], captured on streptavidin-coated magnetic beads (Dynabeads M-280 Streptavidin, Invitrogen). At each round, beads were loaded with 1 µg of biotinylated BMP-1 [catCUB1CUB2] in the presence of the BMP-1 inhibitor UK 383,367 (500 nM) to prevent proteolytic degradation. An antigen-free bead-only condition was processed in parallel as a background control. Beads were blocked for 1 h at room temperature using a different blocking agent at each round (1% casein, 1% BSA and Pierce Protein-Free Blocking Buffer for rounds 1 to 3, respectively) to minimize enrichment of binders specific to blocking agent. The beads were then incubated for 1 h at room temperature with the phage library, which had been pre-blocked in the corresponding blocking agent. Unbound phages were removed by 10 washing cycles with PBS containing 0.1% Tween-20 prior to elution. Bound phages were eluted with 100 mM triethylamine (pH 11.5) and immediately neutralized with 1 M Tris-HCl (pH 8.0). Eluted phages were used to infect exponentially growing *E. coli* TG1 cells and new phage particles were produced following infection with M13K07 helper phage. The phages produced were precipitated from the culture supernatant using PEG/NaCl, quantified by measuring absorbance at 260 nm and used for the subsequent cycle.

After the third panning round, ninety individual TG1 colonies from rounds 2 and 3 were randomly picked using a Hamilton Microlab STARlet colony-picking robot (Robotein®, CIP, ULiège) and grown in 96-deep-well plates for small-scale periplasmic VHH production for 4 h at 37°C. VHH expression was then induced with 1 mM IPTG for 4 h at 37°C. Cytoplasmic extracts, generated by a freezing/thawing cycle were screened by ELISA against BMP-1 [catCUB1CUB2] captured on streptavidin-coated plates, in the presence of the BMP-1 inhibitor as described above. Bound VHHs were detected sequentially with a mouse anti-HA antibody (BioLegend, clone 16B12, 1:2,000) and an alkaline-phosphatase-conjugated goat anti-mouse antibody (Bethyl, A90-116AP, 1:2,000). The signal was developed using pNPP as substrate and changes in absorbance were monitored at 405 nm. Clones showing a specific positive-to-negative signal ratio were selected as candidate binders. The presence of a VHH gene was subsequently verified by colony PCR using MP57/GIII primers pair (Eurogentec), prior to Sanger sequencing. Three non-redundant VHH sequences, designated NbB198, NbB207 and NbB220 were identified and found to belong to three distinct families based on their CDR3 sequences. The corresponding VHH genes, cloned in the pMECS phagemid vector, coding for a C-terminal HA-tag and 6His-Tag, were used to transform *E. coli* WK6 for periplasmic expression and subsequent biochemical and biophysical characterization.

### SPR experiments

Surface plasmon resonance experiments were run on a Biacore T200 apparatus (Cytiva) equipped with the Biacore T200 control software (v3.2.1). Full-length BMP-1 or PCPE-2 were covalently immobilized on Series S CM5 sensor chips (Cytiva) by amine coupling chemistry, using the reagents included in the amine coupling kit (Cytiva). Ligands were diluted in 10 mM sodium acetate pH 4.5 for BMP-1 or in 10 mM HEPES pH 7.4 for PCPE-2. The biotinylated BMP-1 [catCUB1CUB2] was captured on Series S sensor chip SA (Cytiva) following manufacturer’s instructions (Cytiva). The nano-antibodies against BMP-1 were similarly captured on a Series S CM5 sensor chip (Cytiva) after anti-His antibody immobilisation using the His capture kit (Cytiva). SPR signals were recorded simultaneously on a reference flow cell subjected to the same immobilisation/capture procedures except for the presence of protein ligands. Soluble analytes were injected at 50 µL/min at 25°C after dilution in running buffer (10 mM HEPES pH 7.4, 0.15 M NaCl, 5 mM CaCl_2_ and 0.05% P20) for 90 or 120 s. Regeneration was achieved with 2 M guanidium chloride for the CM5 chip and with 10 mM glycine pH 1.5 for the anti-His capture. Kinetic and steady-state analyses were carried out using the Biacore T200 Evaluation software (v3.2.1).

### Cleavage assays

All cleavage assays were performed with 13 nM BMP-1 at 37°C in 50 mM HEPES (pH 7.4), 5 mM CaCl_2_ and 0.02% βOG. The concentration of the recombinant ectodomain of human LDLR (Bio-techne), PCPE-2 and deletion mutants was 388 nM. Cleavage products were analysed by SDS-PAGE using 4-20% Criterion polyacrylamide gels (Bio-Rad) followed by staining with InstantBlue (Euromedex). Quantification of protein band intensities was made with the ImageQuant TL software v8.2 (Cytiva). Cleavage of the fluorogenic peptide Mca-YVADAPK(Dnp)-OH (20 μM, Covalab) was as described previously^49^. Fluorescence was measured for 15 min (excitation: 320 nm; emission: 390 nm) using a Tecan Spark® fluorimeter controlled by the SparkControl software (v3.2) and the slope of the curve was used to derive BMP-1 activity.

### HDX-MS

For peptide mapping, BMP-1 [catCUB1CUB2] diluted in a non-deuterated buffer was subjected to the same digestion protocol and liquid chromatographic gradient as detailed below for the deuterated samples. MSE analysis was performed with a Synapt XS mass spectrometer (Waters), applying collision energy ramping from 20 to 30 kV. Sodium iodide was used for calibration and leucine enkephalin was applied for mass-accuracy correction. MSE runs were analysed with ProteinLynx Global Server (PLGS) 3.0 (Waters), and peptides identified in 3 out of 4 runs, with at least 0.2 fragments per amino acid and at least 2 fragments in total and 1 consecutive fragments, and with mass error below 7 ppm, were selected in DynamX 3.0 (Waters). For deuterium labelling, 10 µM of BMP-1 and 10 µM of BMP-1 in complex with PCPE-2 CUB1CUB2 were diluted 1:10 in a fully deuterated labelling buffer at 20 mM HEPES, 250 mM NaCl, 5 mM CaCl_2_, 0.1% DDM, (pD = 7.3 at 23 °C). The HDX reactions were conducted for 8 s, 2 min, 30 min and 450 min at 23 °C, and quenched by a 1:1 dilution (vol/vol) with an ice-cold 100 mM phosphate buffer containing 4 M Urea and 100 mM TCEP (pHread=2.3). Samples were held for 30 s on ice and snap-frozen in liquid nitrogen. Frozen samples were quickly thawed and injected into an Acquity UPLC M-Class System with HDX Technology (Waters). Injected samples first passed at 110 µL/min through a home-made pepsin column and a PNGaseRc column (AffiPro) at 20°C for on-line digestion and N-glycan removal, respectively. Peptides were trapped/desalted with solvent A (0.23% formic acid in water, pH 2.5) for 4 min at 110 μL/min and at 0°C through an Acquity BEH C18 VanGuard pre-column (1.7 μm, 2.1 mm × 10 mm, Waters). Peptides were eluted into an Acquity UPLC BEH C18 analytical column (1.7 μm, 2.1 mm × 50 mm, Waters) with a 7-min linear gradient rising from 8% to 35% solvent B (0.23% formic acid in acetonitrile) at a flow rate of 90 μL/min, at 0°C. Peptides were then subjected to electrospray ionization in positive mode and MS analysis with ion-mobility separation. Triplicates were conducted at time points 2 min and 450 min, duplicates were conducted at time points 8 s and 30 min. Peptide-level deuterium uptake was calculated with DynamX 3.0, after data were visually inspected and curated. The threshold for statistically significant differences in HDX was established at the 98% confidence level, based on an approach described previously^50^. Overlapping peptides were used to define the areas of HDX changes. As per community-based recommendations^51^, a summary of the HDX-MS experiments is provided in Table S3.

### Cryo-EM

To obtain the structure of the PCPE-2/BMP-1 complex, 2 nmol of each purified protein were mixed with 250 mM of Methyl-PEG4-NHS-Ester (Thermo Fisher Scientific) and incubated on ice during 1 h. The pegylation reaction was quenched by the addition of 0.1 M of Tris pH 7.5. The pegylated complex was purified by SEC on a Superdex 200 pg 10/300 column in 20 mM HEPES pH 7.4, 0.15 M NaCl, 2.5 mM CaCl_2_.

Grids were prepared using a Vitrobot Mark IV plunge freezer (FEI) at 100 % humidity and 4°C. UltrAuFoil (Quantifoil) (R2/2 on a 200 mesh) were glow discharged, before a volume of 3.5 µL of sample at 1.2 mg/mL was applied and blotted for 3.5 sec before vitrification in liquid ethane. Cryo-EM data were collected at the Oxford Particle Imaging Centre, on a 300 kV G3i Titan Krios microscope (Thermo Fisher Scientific) equipped with a SelectriX energy filter and Falcon IV direct electron detector. Data were collected automatically using EPU and a 30 degrees stage tilt. Movies were recorded in EER format with a total dose of -55e/Å^2^ and a calibrated pixel size of 0.73 Å /px.

Data were processed using cryoSPARC^52^. EER format movies were fractionated in 55 frames. Patch motion correction and patch CTF-estimation were performed with default setting. Corrected micrographs with poor statistics were manually curated. Micrograph denoiser and junk detector were used to improve particle picking. First, particles of interest were picked using blob and template picker were subjected to multiple rounds of 2D classification. After 2D classification, well-resolved classes were selected, and three ab initio models were generated and further refined using heterogeneous refinement. Particles belonging to the complex class were used to train a Topaz model and pick a new set of particles. These particles were directly classified using heterogeneous refinement with the previous ab initio models input. One good 3D class containing 170,000 particles and representing the complex was selected and refined using non-uniform refinement. Additionally, to limit the impact of preferential orientation/flexibility homogenous ab-initio refinement followed by local refinement was used, leading to a final reconstruction at 3.64 Å nominal resolution.

#### Model building and refinement

AlphaFold3 prediction was performed using AlphaFold3 web server (https://alphafoldserver.com)^31^. The sequences of PCPE-2 and BMP-1, as documented in Table S2, were used, with the addition of one zinc ion and three calcium ions. Three different seeds (seed numbers chosen automatically) were used to calculate 15 final models (5 per job) which were all analysed, and the highest confidence model was finally used.

CryoZeta was used to create an accurate structural prediction consistent with the Cryo-EM density map. The software computes a residue-level confidence score by combining local map-to-model cross correlation with structural confidence metrics, allowing the identification of regions that are well supported by the experimental density. The FASTA sequences of BMP-1 [catCUB1CUB2] and PCPE-2 CUB1CUB2 and the corresponding cryo-EM map were provided to the CryoZeta server (https://em.kiharalab.org/algorithm/CryoZeta), using a contour level of 0.0607 and a resolution of 6 Å. Although the nominal resolution of the cryo-EM reconstruction was estimated at 3.64 Å, value of 6 Å was used for CryoZeta modelling as it more accurately reflected the level of structural detail observed in the density map. The model reliability score, that permit to compare the models, was calculated by using this formula:

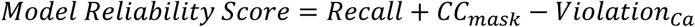

where Recall represents the fraction of predicted Cα/C1’ atoms with at least one support point within 3 Å; CC corresponds to the cross-correlation coefficient between the density simulated from the predicted structure and the experimental cryo-EM map, calculated over voxels within a defined region surrounding the molecular structure; and Violation denotes the number of Cα-Cα distances between consecutive residues exceeding 4 Å.

#### Interface analysis

Protein-protein interfaces were analysed using the PDBePISA server^53,54^ (Protein Interfaces, Surfaces and Assemblies) to identify and characterize the intermolecular contacts within CUB2-CUB2 and CUB1-catalytic domain in the complex. Only the buried surface area was analysed.

Molecular models were visualized and structurally aligned using PyMol (The PyMol Molecular Graphics System, Version 4.6, Schrödinger, LLC). RMSD values were calculated in PyMol to assess the structural similarity between the models.

## Statistical analysis

Statistical analyses were performed using GraphPad Prism version 10. Data are presented as mean ± standard deviation (SD). Comparisons between multiple groups were performed using One-way ANOVA, followed by Dunnett’s multiple comparison test with each condition compared with the BMP-1 control.

## Supplementary data

**Figure S1:**
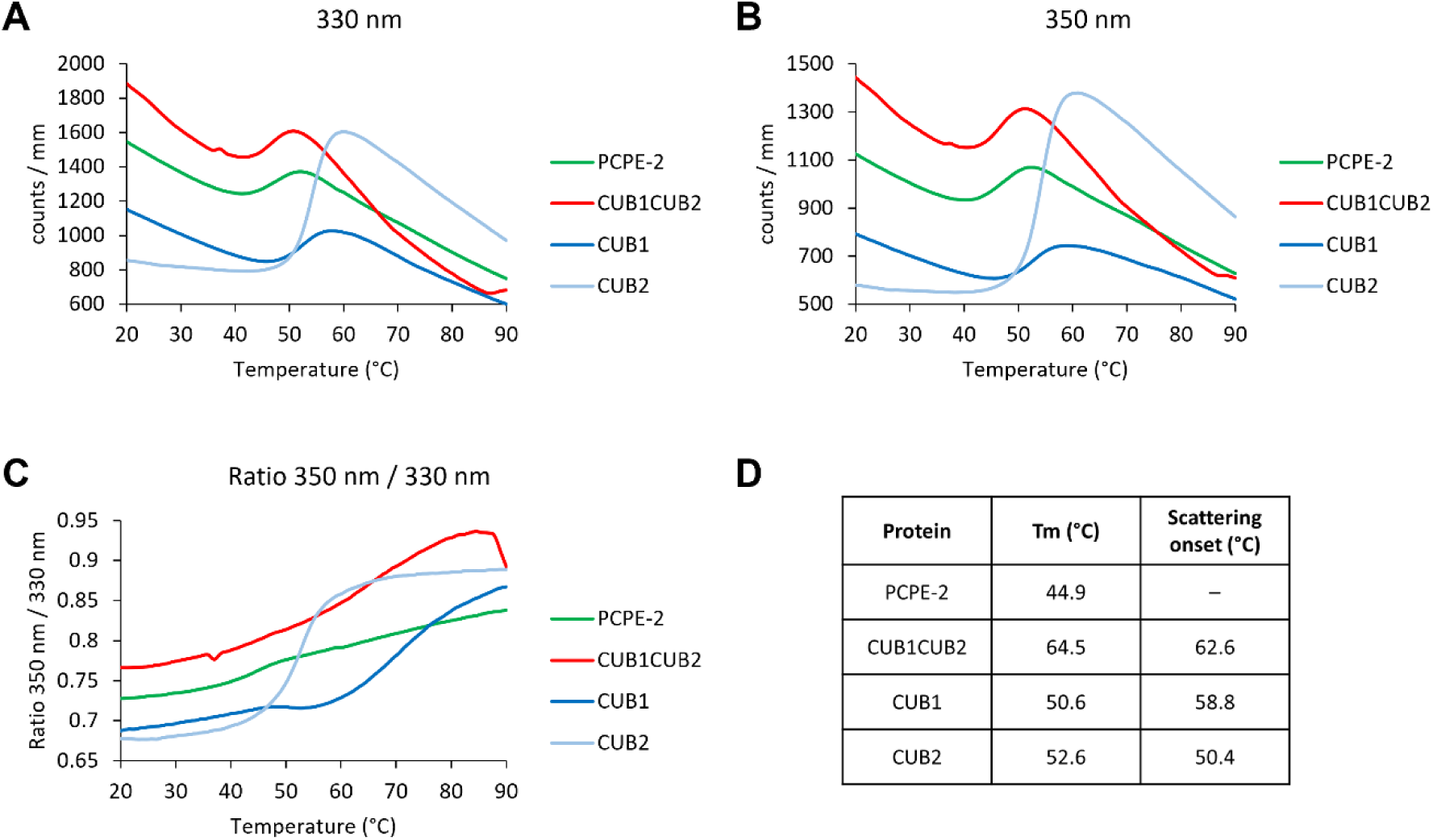
Thermal stability profiles of PCPE-2 constructs assessed by nanoDSF. Changes in fluorescence intensity at 330 nm **(A),** at 350 nm **(B)**, and ratio 350 nm / 330 nm **(C)** as a function of temperature for PCPE-2 and its domains. **D.** Tm values and scattering onset temperatures, calculated from the 350nm/330nm first derivative.

**Figure S2:**
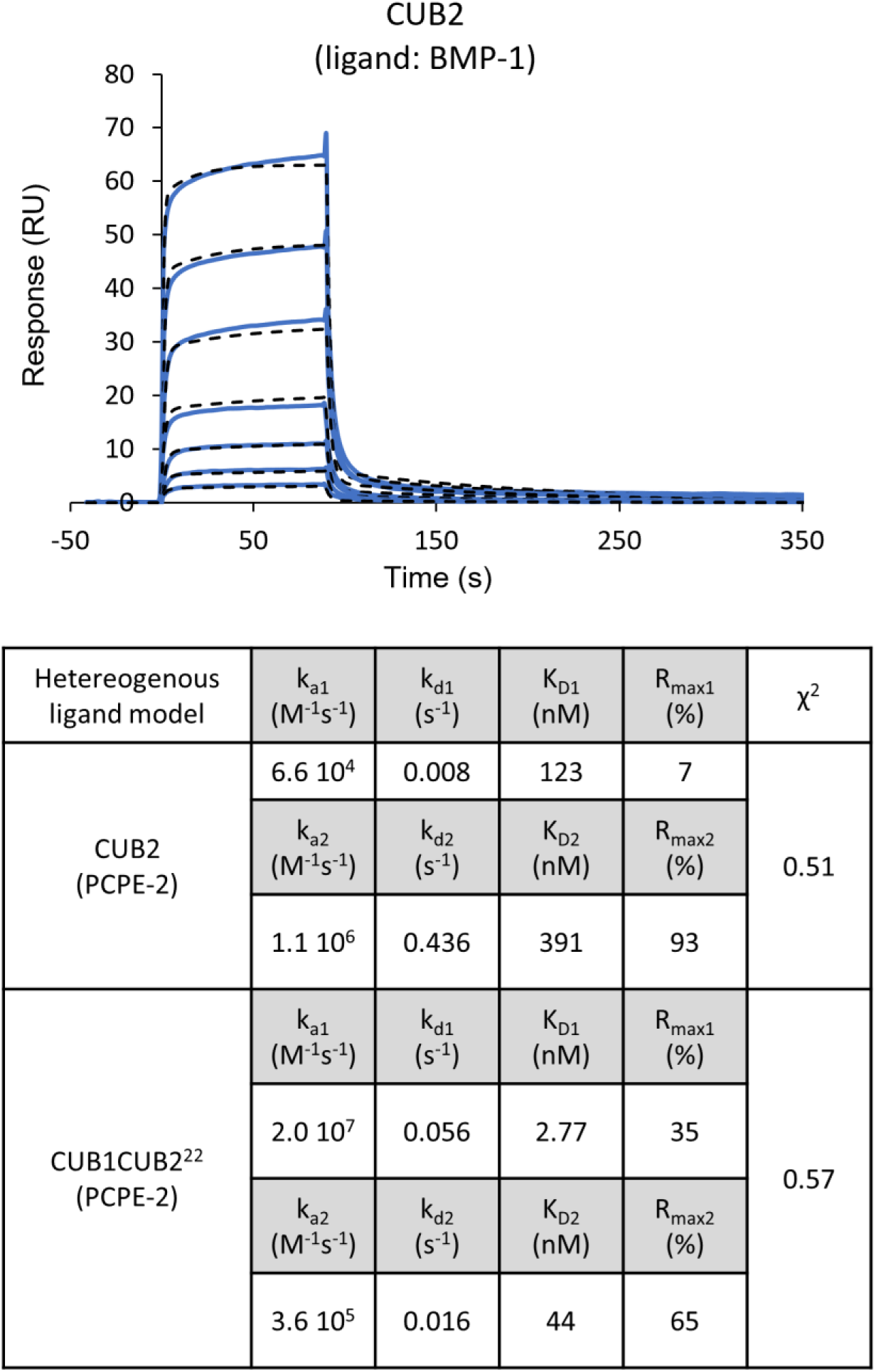
Kinetics parameters of CUB2 (PCPE-2). CUB2 binding to immobilized BMP-1 (1640 RU). Increasing concentrations of PCPE-2 were injected (12.5 – 800 nM, prepared as serial two-fold dilutions). Fits were obtained with the kinetic model (black dotted lines; heterogeneous ligand). For comparison, values obtained for CUB1CUB2 (extracted from^22^) are also indicated.

**Figure S3:**
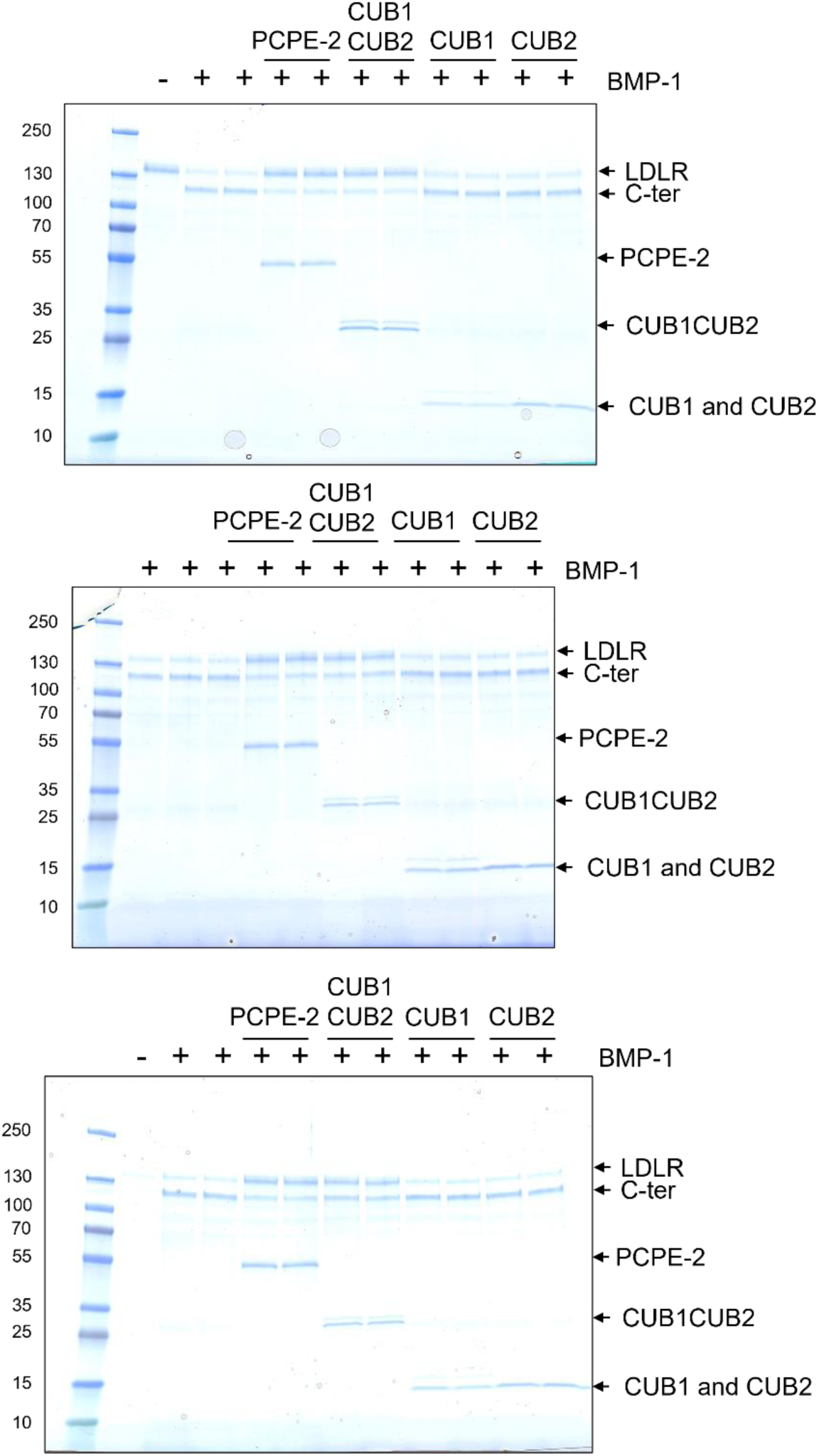
Uncropped gels of LDLR activity assays. Cleavage of LDLR ectodomain (388 nM) by BMP-1 (13 nM) in the presence of PCPE-2 or its deletion mutants (388 nM). C-ter = LDLR C-terminal fragment after BMP-1 cleavage.

**Figure S4:**
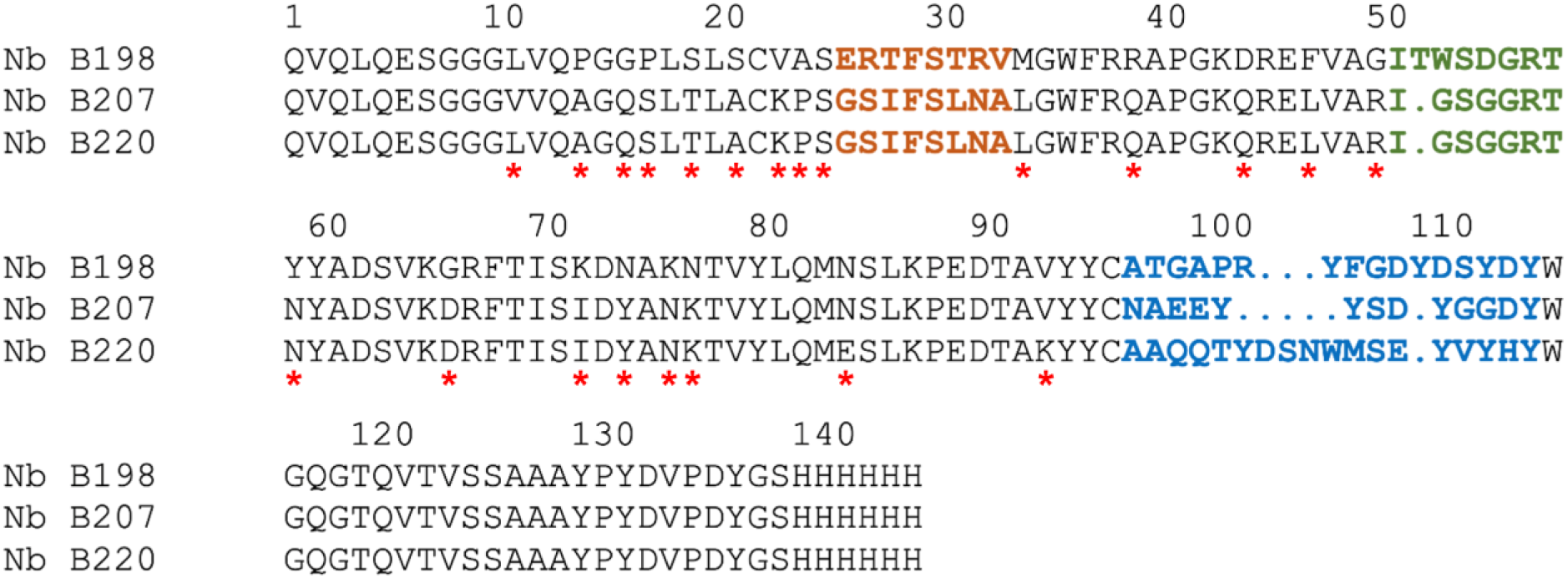
Alignment of the three anti-BMP-1 VHHs. The CDR1, CDR2 and CDR3 of each VHH are indicated respectively in orange, green and blue. Red stars indicate amino acid substitutions in the framework.

**Figure S5:**
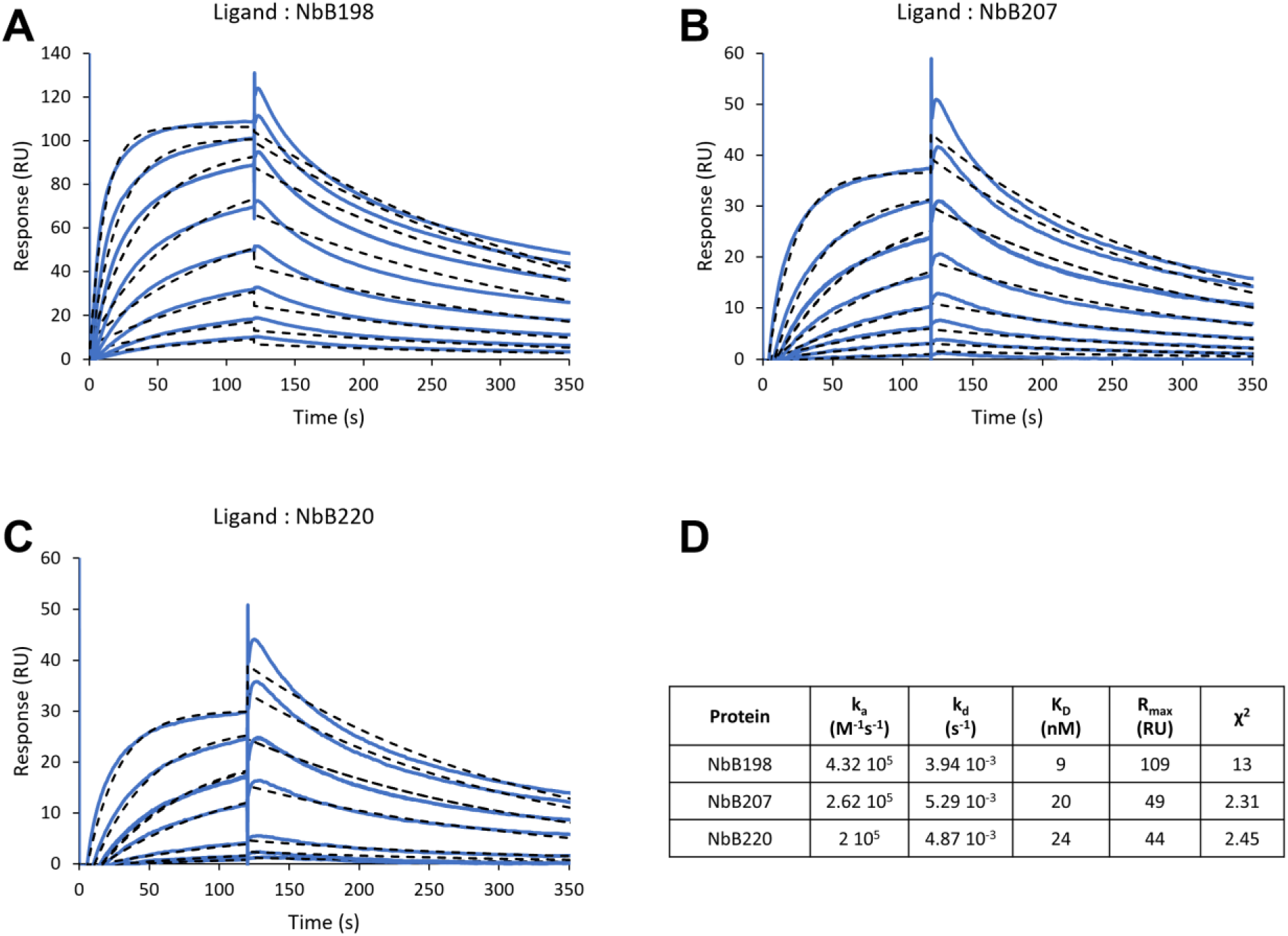
Analysis of the interaction of VHHs with BMP-1. **A.** Fit (1:1 model) of sensorgrams obtained when increasing concentrations of BMP-1 (1.56 nM – 200 nM) were injected over immobilized NbB198 (34 RU). **B.** Same as (A) with NbB207 (17 RU). **C** Same as (A) with NbB220 (12 RU). **D.** Table of the kinetic data obtained from the fits.

**Figure S6:**
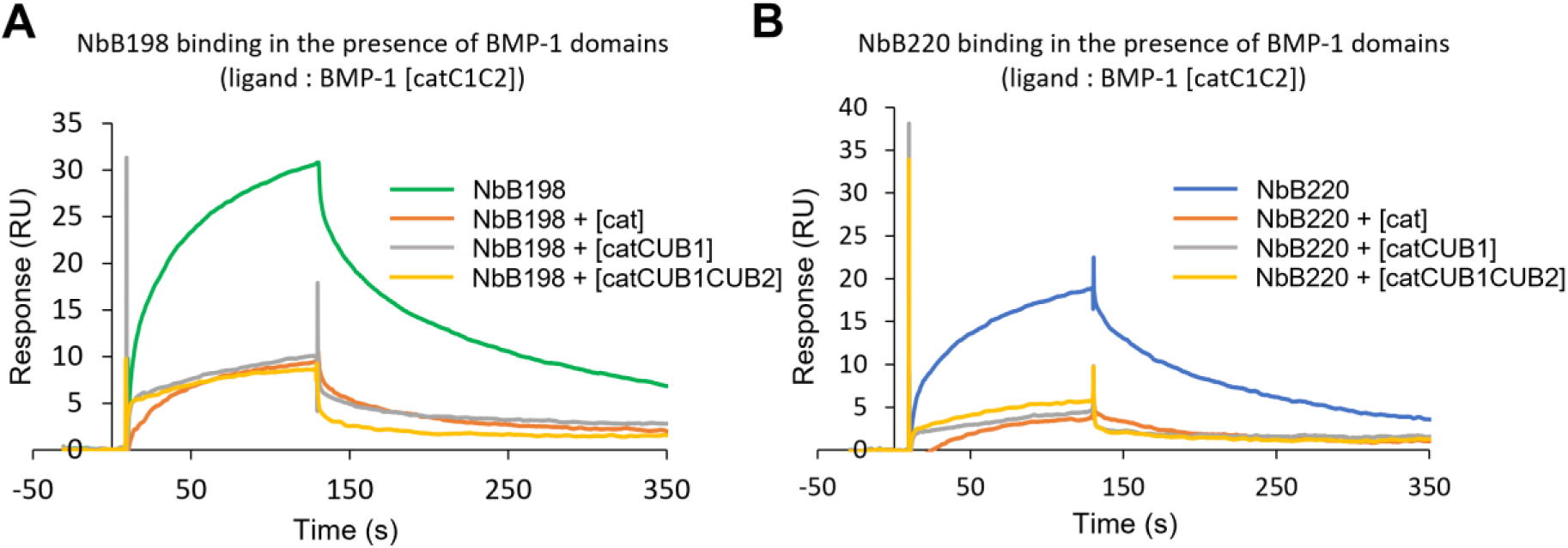
NbB198 and NbB220 also interact with the catalytic domain of BMP-1. **A.** Competition experiment in which 100 nM of NbB198 was co-injected with 100 nM of BMP-1 truncated mutants ([cat], [catCUB1], [catCUB1CUB2]). The immobilized protein was [cat-CUB1CUB2] BMP-1 (397 RU). **B.** Same as (A) for NbB220.

**Figure S7:**
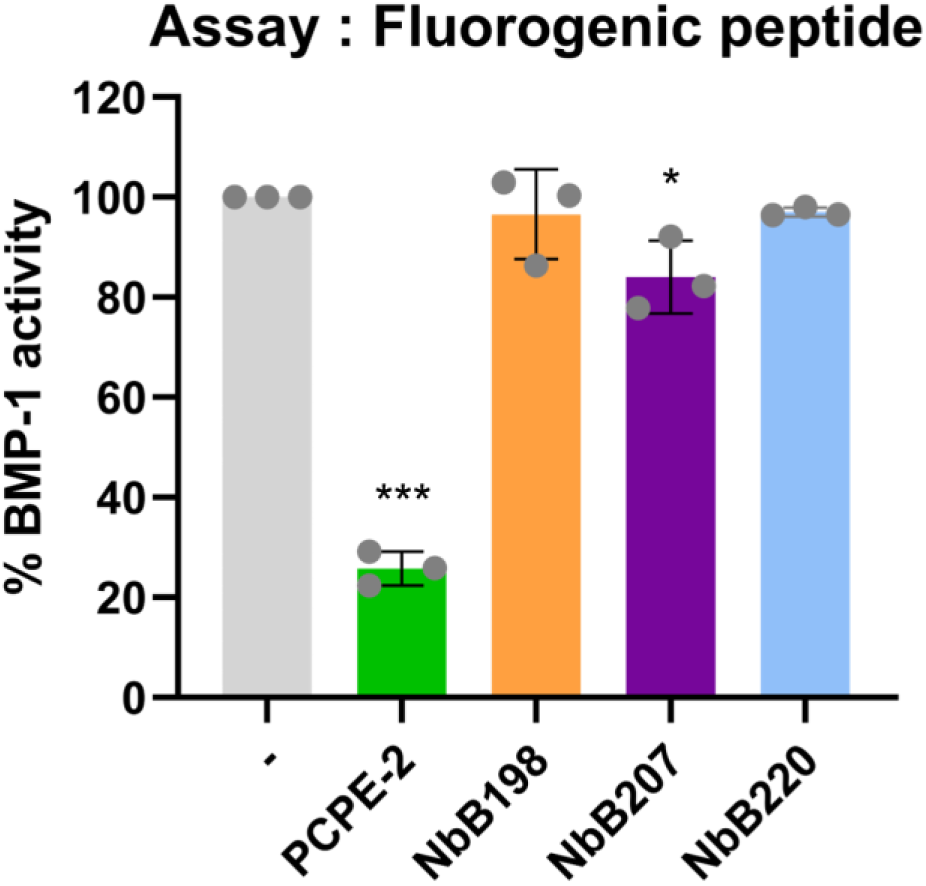
Effect of the VHHs on BMP-1 activity. Quantification of BMP-1 activity with the Mca-YVADAPK(Dnp)OH fluorogenic peptide in the presence of 1 µM of VHHs. BMP-1 concentration in the experiment was 7 nM and PCPE-2 was 50 nM. Means ± SD of n = 3 independent experiments performed in duplicate. Statistical significance (comparison with the BMP-1 alone condition): p-value ≤ 0.05 (*) and p-value ≤ 0.001 (***).

**Figure S8:**
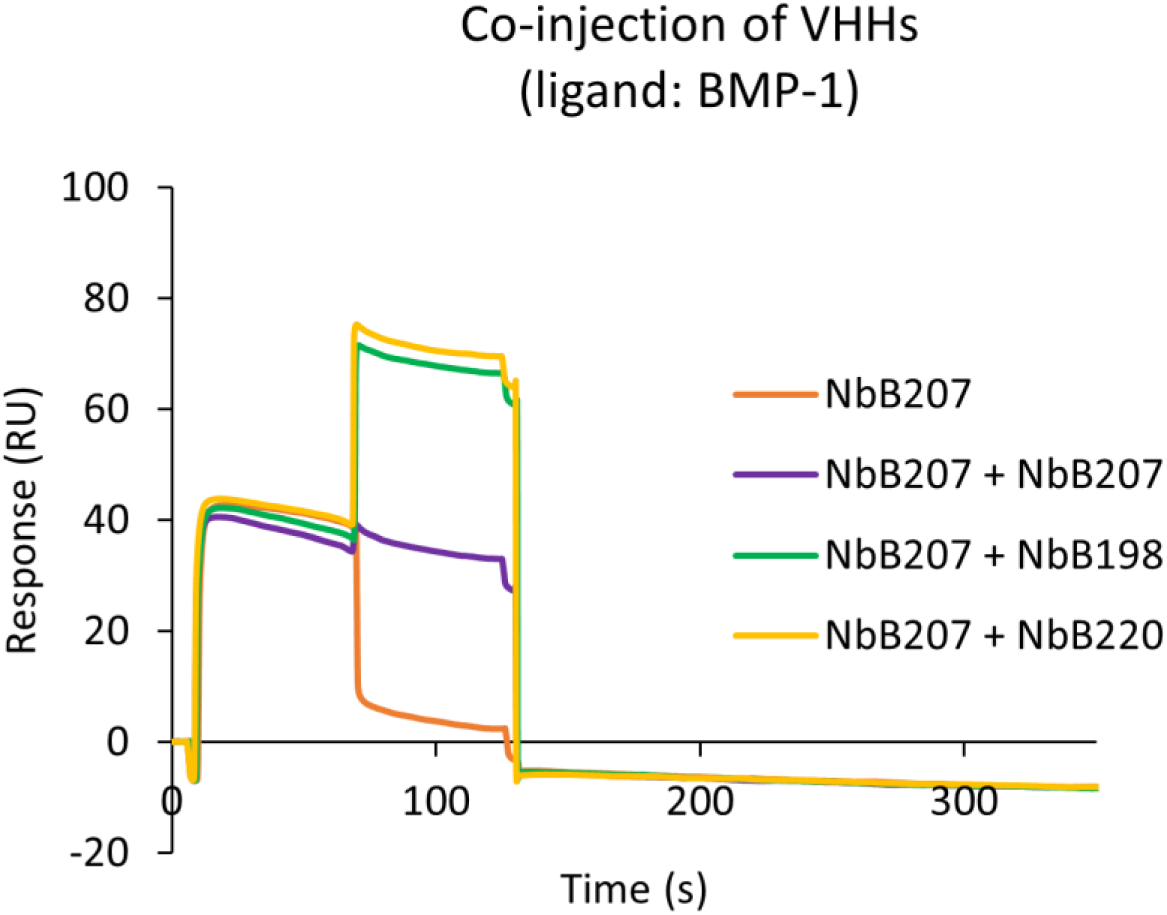
Competition experiments consisting of an injection of 1 µM NbB207 for 60 s followed by an injection of NbB207 (1 µM) alone or in combination with 1 µM NbB198 or NbB220 for another 60 s (dual inject mode). The immobilized protein was BMP-1 (1640 RU).

**Figure S9:**
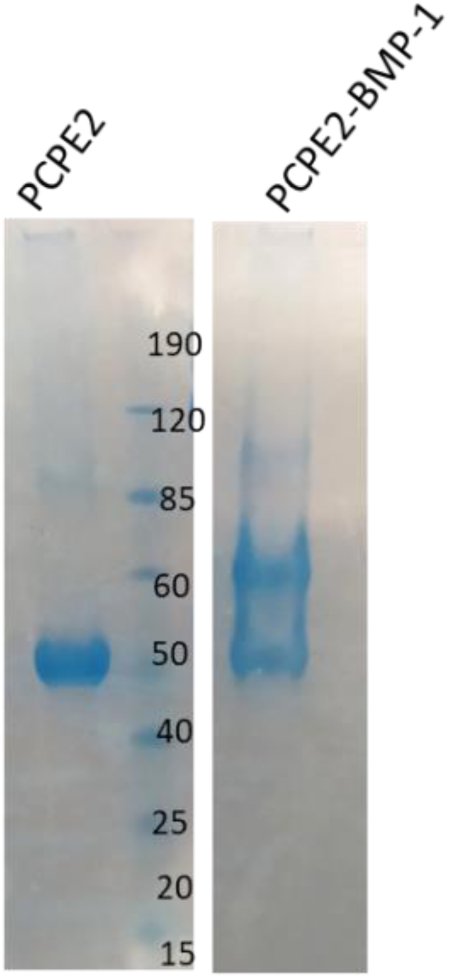
SDS-page analysis of BMP-1/PCPE-2 complex after PEGylation and purification (non-reducing conditions).

**Figure S11:**
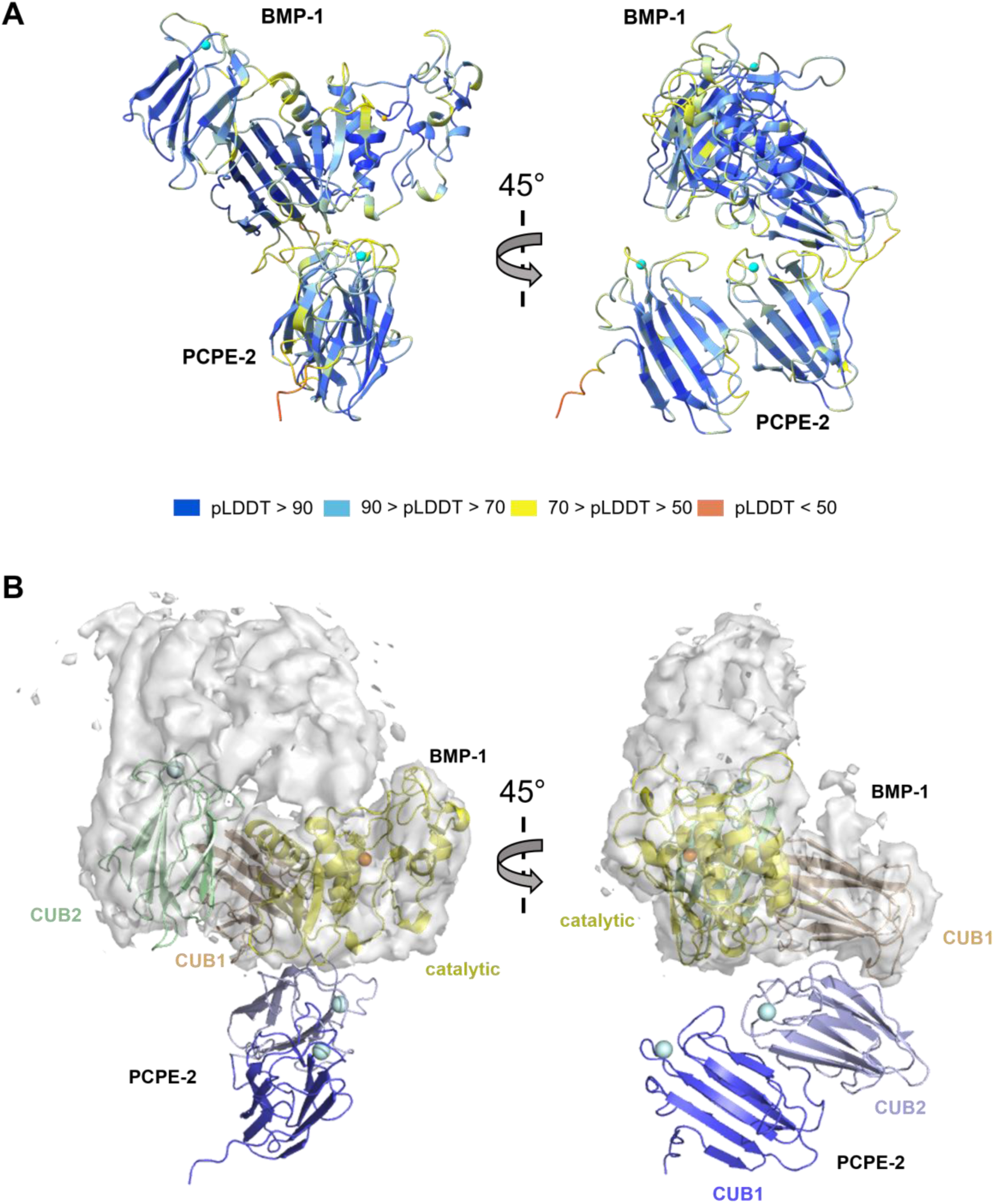
Analysis of the PCPE-2/BMP-1 AlphaFold3 model and placement inside the cryo-EM mesh. **A.** pLDDT scores of the AlphaFold model of the BMP-1 [catCUB1CUB2] – PCPE-2 CUB1CUB2 complex. The chains of each protein are coloured by the pLDDT scores. **B.** Placement of the AlphaFold3 model inside the cryo-EM density map based on the optimal position for BMP-1.

**Figure S12:**
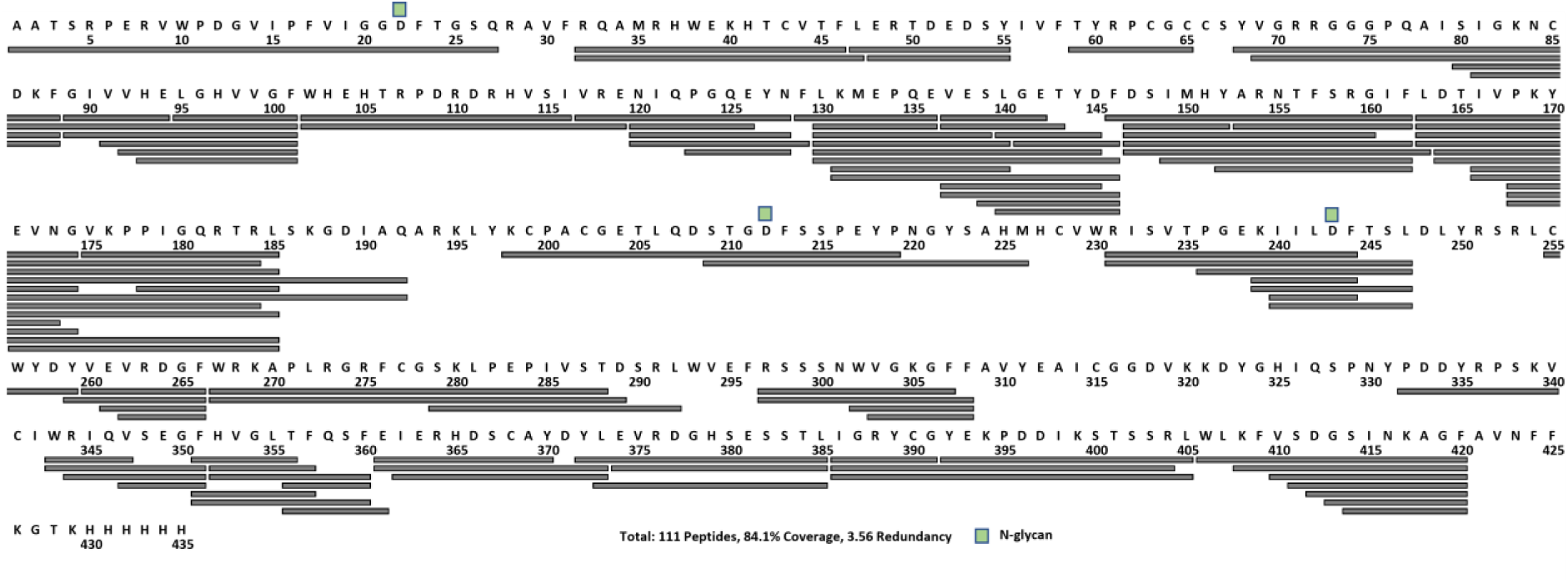
Peptides of BMP-1 whose HDX was followed are depicted with gray bars along the protein sequence. Green squares indicate N-glycosylation sites, while the amino acid D shown below each square corresponds to a deaminated asparagine residue.

**Figure S13:**
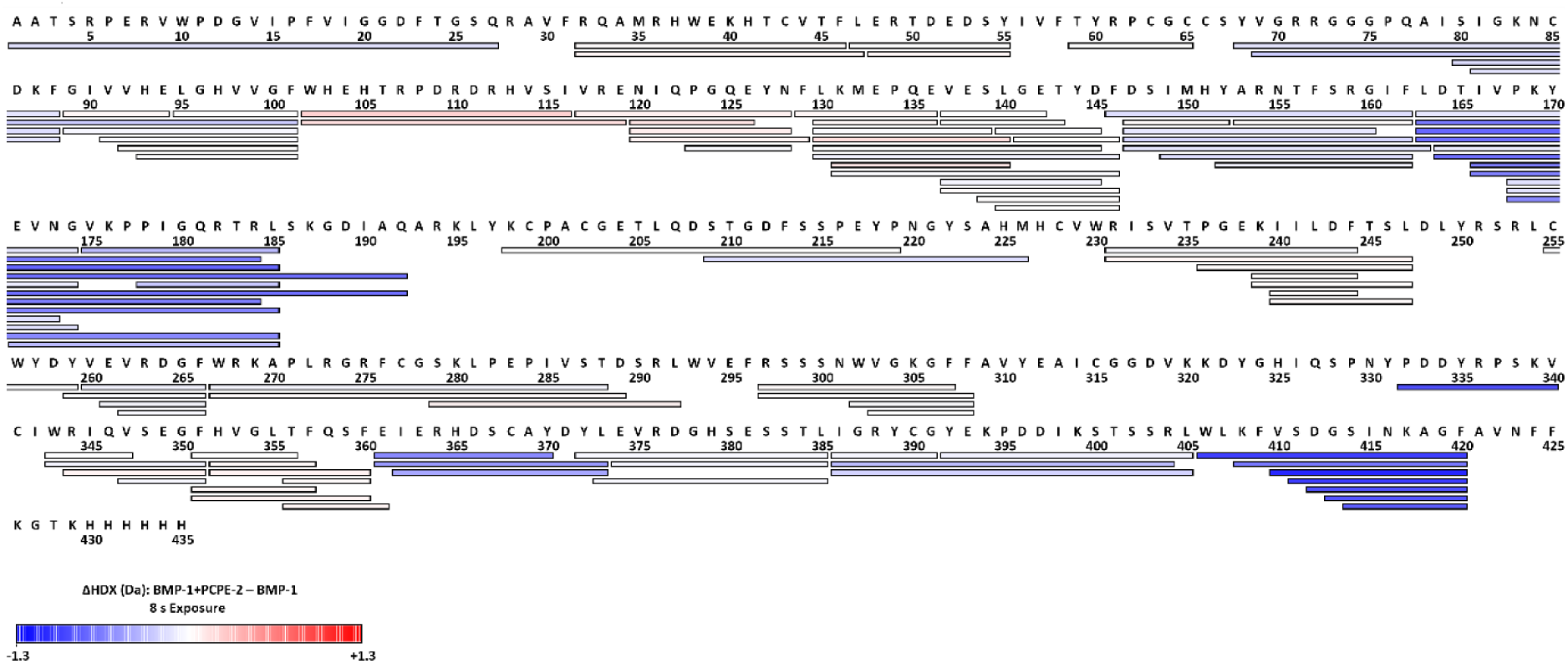
Differences in HDX (ΔHDX, Da) at 8 s are color-coded along the peptides studied.

**Figure S14:**
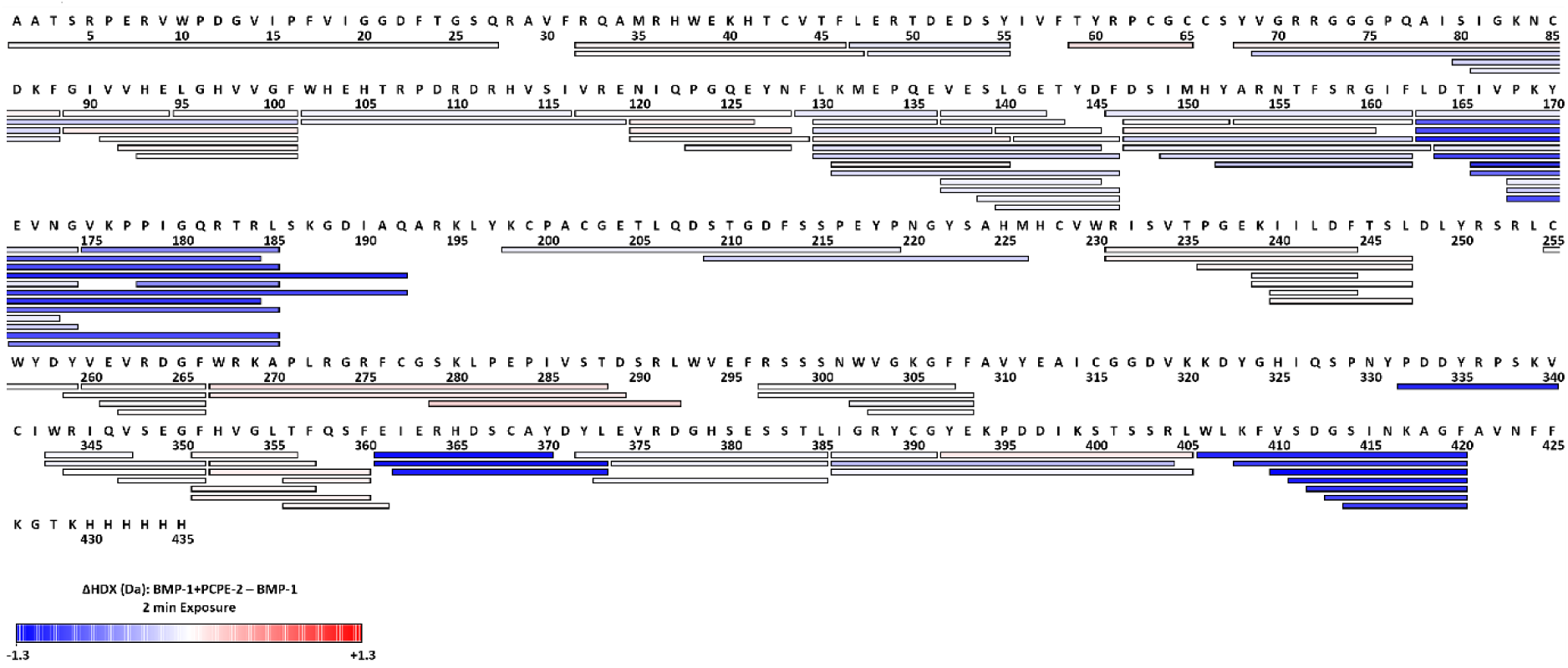
Differences in HDX (ΔHDX, Da) at 2 min are color-coded along the peptides studied.

**Figure S15:**
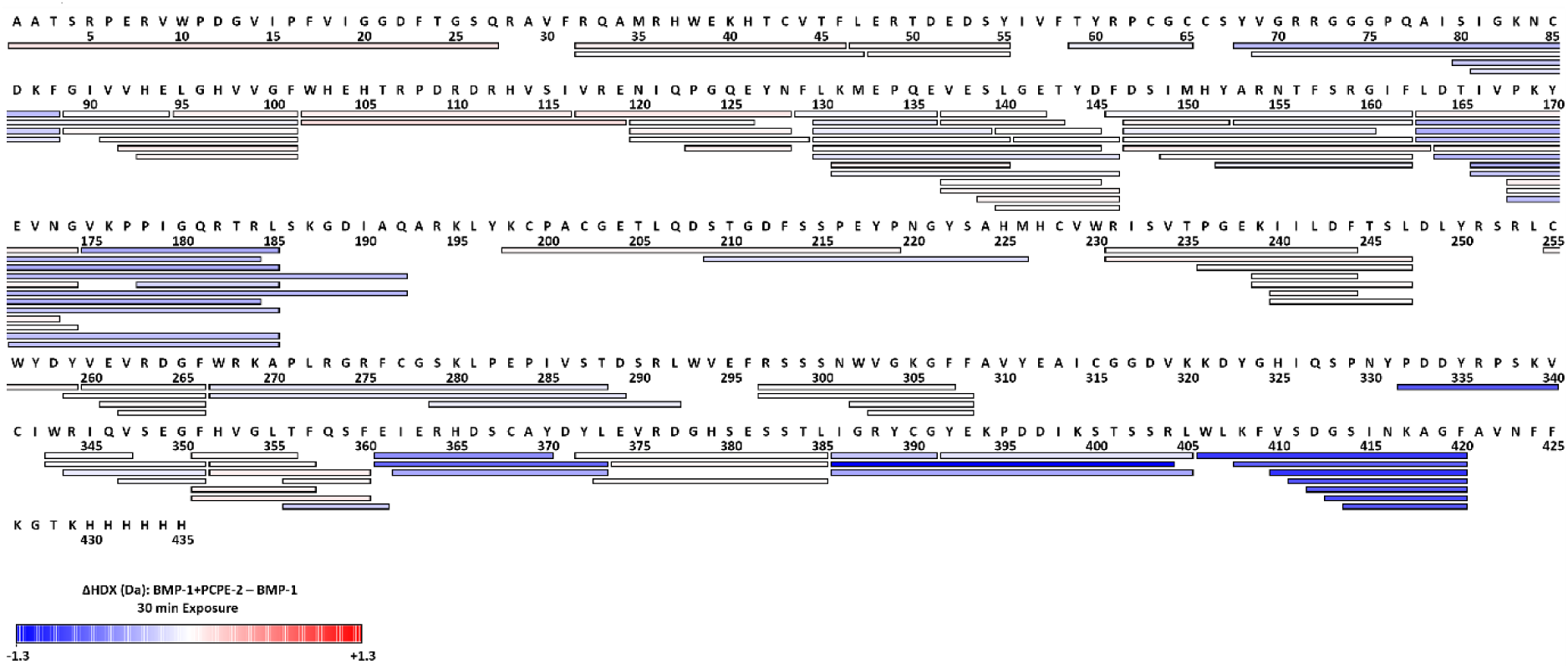
Differences in HDX (ΔHDX, Da) at 30 min are color-coded along the peptides studied.

**Figure S16:**
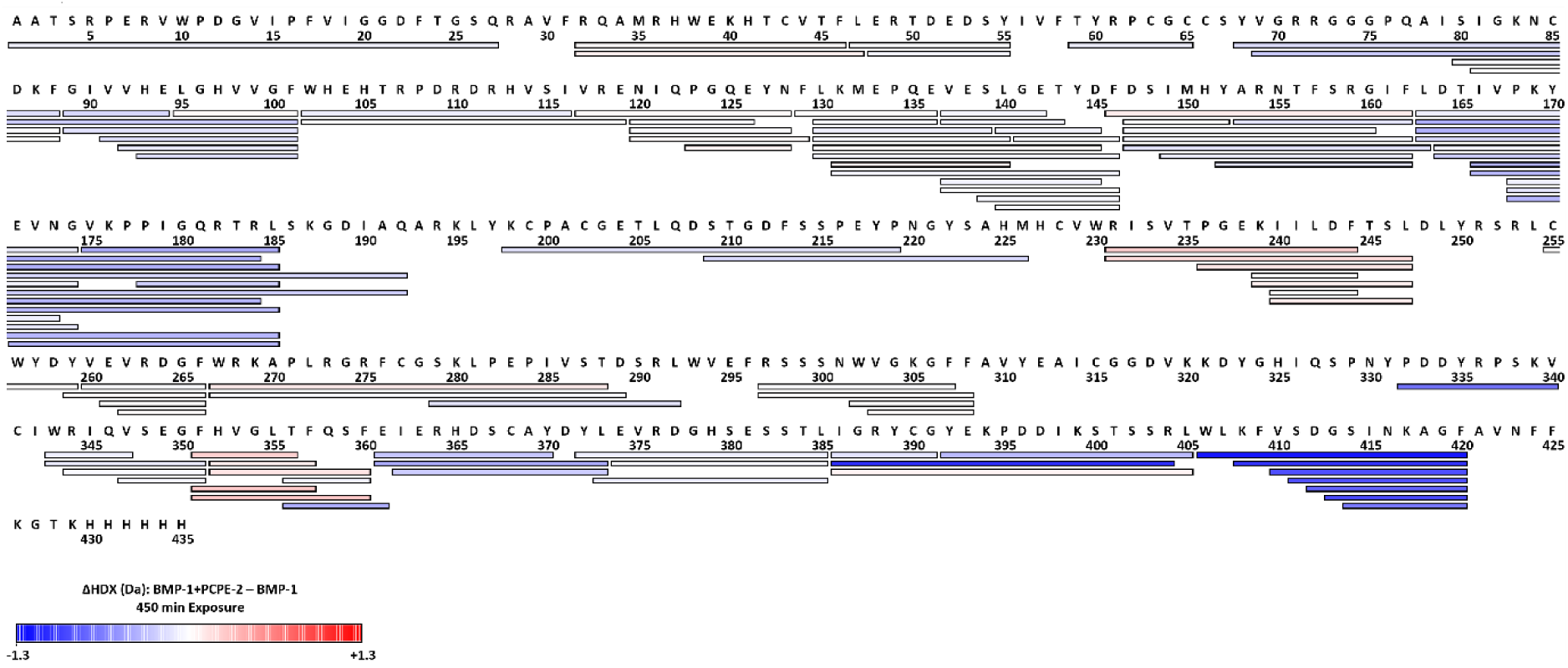
Differences in HDX (ΔHDX, Da) at 450 min are color-coded along the peptides studied.

**Table S1:** HDX summary table.

| HDX Data Set | Protein alone vs complex |
| --- | --- |
| Labelling buffer | 20 mM HEPES, 250 mM NaCl, 5 mM CaCl<sub>2</sub>, 0.1% DDM (pD = 7.3), D<sub>2</sub>O% = 90% |
| HDX time course | 8 s, 2 min, 30 min, 450 min (at 23 °C) |
| HDX control samples | Maximally-labelled control not performed |
| Back-exchange (mean / IQR) | N/A |
| # of Peptides | 111 |
| Sequence coverage | 84.1% |
| Average peptide length / Redundancy | 11.75 / 3.56 |
| PTMs detected | N-glycan at N22, N212, N243 |
| Replicates (technical) | triplicates at time points 2 min and 450 min; duplicates at time points 8 s and 30 min |
| Repeatability (average standard deviation) | protein alone: 0.062922; complex: 0.043032 |
| Significant differences in HDX (delta HDX > X D) | 0.42 Da (98% CI) |

**Table S2:**
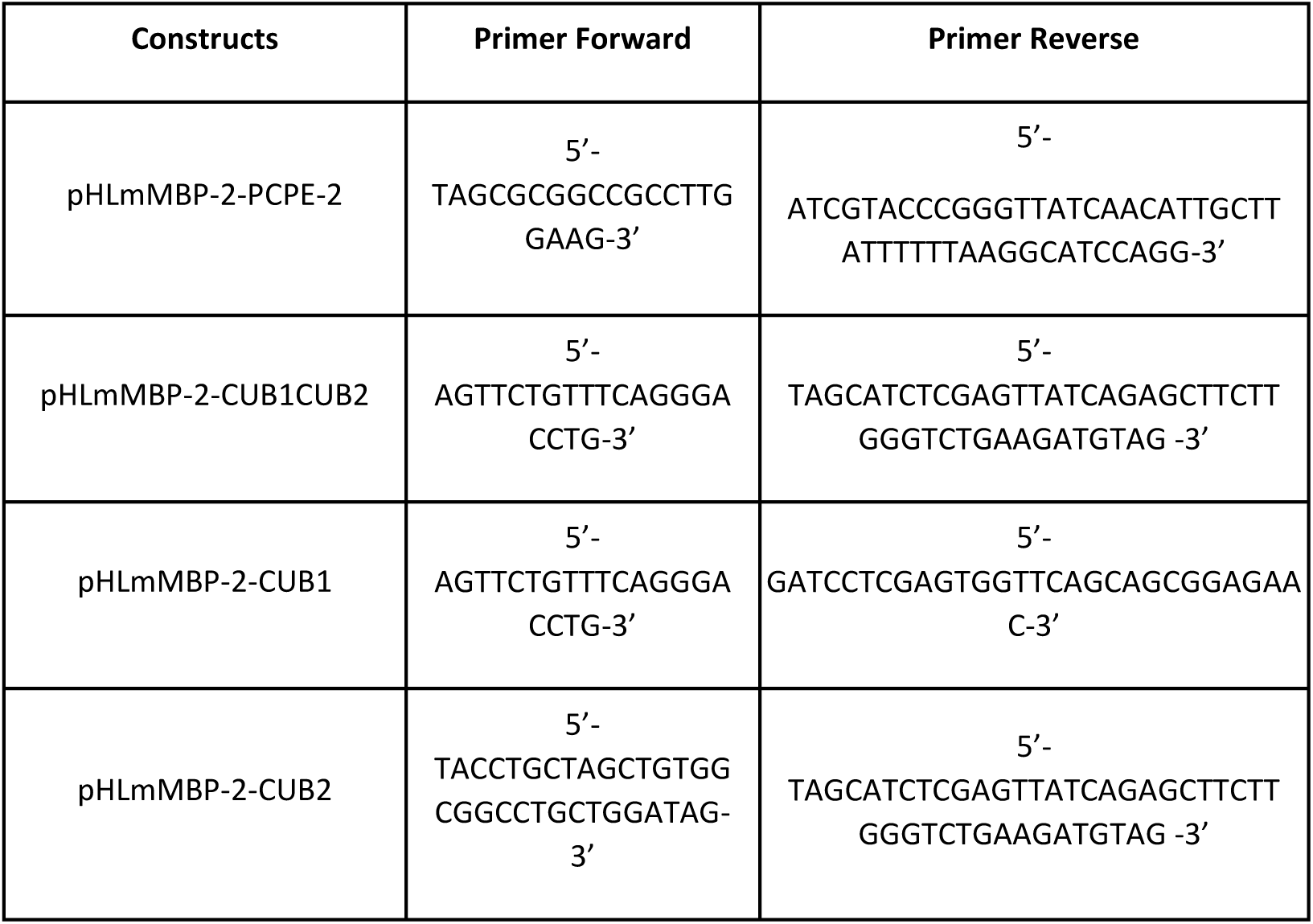
Primers used for PCR experiments.

**Table S3:**
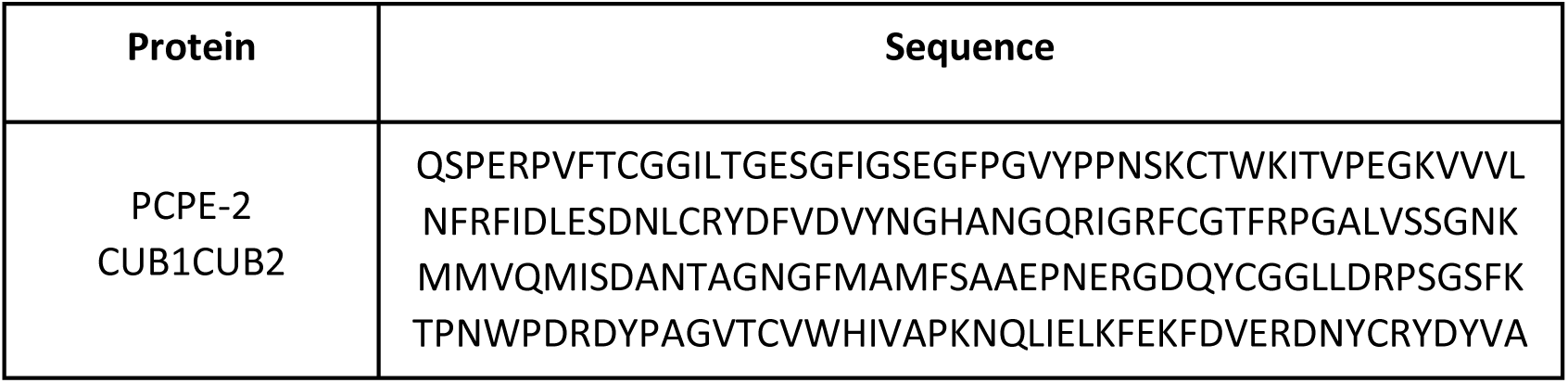
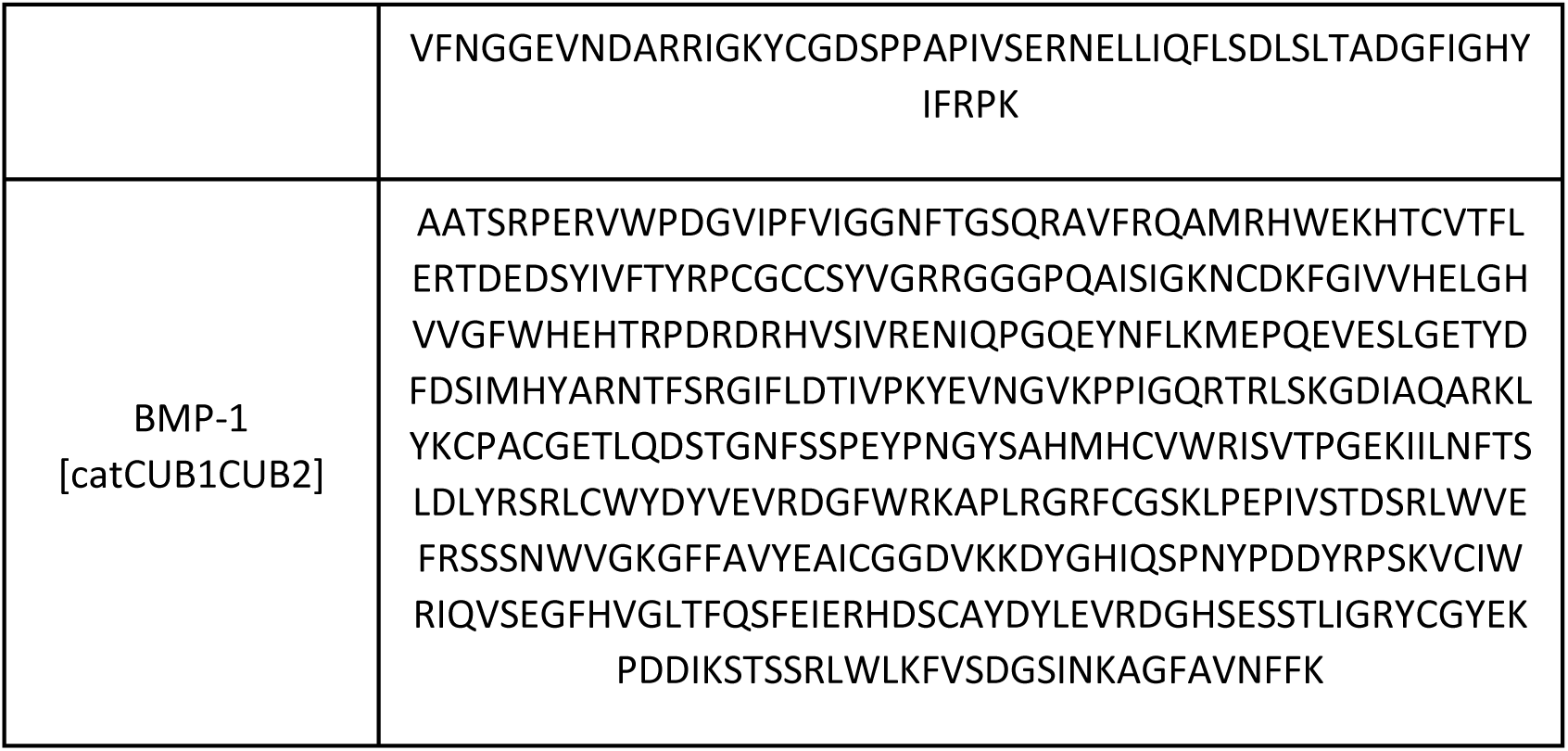
Sequences of PCPE-2 CUB1CUB2 and BMP-1 [catCUB1CUB2] (mature forms).

## Acknowledgements

We would like to thank Cindy Dieryckx and Camille Callies for their help in protein production, and Oskar Lipińsky for his assistance in grid preparation training and for insightful discussions. We acknowledge the microscopy facility “Centre Technologique des Microstructures CTµ” of Université Lyon 1, member of the national infrastructure France-BioImaging (https://ror.org/01y7vt929) supported by the French National Research Agency (ANR-24-INBS-0005 FBI BIOGEN), for the access to the cryo-EM grid preparation equipment. We also thank the UMS3444 (SFR Biosciences, Lyon) for the access to the Biacore instrument, and Céline Freton from MMSB (UMR 5086, Lyon) for providing access to the Prometheus NanoDSF apparatus. This work benefited from access to the Instruct-ERIC centers Oxford HDX-MS facility (PID34935, VID62893) and Oxford Particle Imaging Centre (OPIC) (PID34935, VID62894). We acknowledge Alain Vanderplasschen for the immunization of the alpaca and the preparation of the PBMCs. We acknowledge the Robotein® platform of the BE Instruct-ERIC Centre for providing access to the EasyPick Microlab STARlet Hamilton workstation (https://www.robotein.uliege.be/cms/c_14301428/en/robotein). This work was performed using the computing facilities of the PRABI Lyon-Gerland.

## Funding

This work was funded by a grant from the Agence Nationale de la Recherche (ANR-21-CE11-0020-01), the CNRS, the Université Lyon 1, Instruct-ERIC (PID34935) and the EDISS doctoral school.

## Notes

### Competing Interest Statement

The authors have declared no competing interest.

